# Hippocampal place cells encode local surface texture boundaries

**DOI:** 10.1101/764282

**Authors:** Chia-Hsuan Wang, Joseph D. Monaco, James J. Knierim

## Abstract

The cognitive map is often assumed to be a Euclidean map that isometrically represents the real world (i.e. the Euclidean distance between any two locations in the physical world should be preserved on the cognitive map). However, accumulating evidence suggests that environmental boundaries can distort the mental representations of a physical space. For example, the distance between two locations can be remembered as longer than the true physical distance if the locations are separated by a boundary. While this overestimation is observed under different experimental conditions, even when the boundary is formed by flat surface cues, its physiological basis is not well understood. We examined the neural representation of flat surface cue boundaries, and of the space segregated by these boundaries, by recording place cell activity from dorsal CA1 and CA3 while rats foraged on a circular track or square platform with inhomogeneous surface textures. About 40% of the place field edges concentrated near the surface cue boundaries on the circular track (significantly above the chance level 33%). Similarly, the place field edges were more prevalent near the boundaries on the platforms than expected by chance. In both 1-dimensional and 2-dimensional environments, the population vectors of place cell activity changed more abruptly with distance between locations that crossed cue boundaries than between locations within a bounded region. These results show that the locations of surface boundaries were evident as enhanced decorrelations of the neural representations of locations to either side of the boundaries. This enhancement might underlie the cognitive phenomenon of overestimation of distances across boundaries.

## Introduction

Real-world space has a universal metric (at least on the local scale of everyday experience), with distance varying regularly along each of 3 dimensions (i.e., a meter measured at each location along the x dimension is equal to a meter along the y and z dimensions). Psychological space, however, can be much more complex [1–3]. For example, compartmentalization of an environment can result in perceptual distortions of the Euclidean space [4–9], such as increasing the mental distance between two locations separated by a boundary [7–9].

The physiological mechanisms underlying such distorted representations of space are not well understood. Tolman suggested that an internal representation of the environment—a “cognitive map”—is used by an organism to devise flexible solutions to various cognitive tasks [10]. The subsequent discoveries of place cells [11, 12], grid cells [13], head direction cells [14, 15] and boundary cells [16–18] in rodents and primates [19–23] provided strong evidence that this map is instantiated in the hippocampus and related structures. The map is generated by an interaction between two major types of neural computation: path integration, the integration of a velocity vector over time to continually update a position estimate based on self-motion, and landmark navigation, the use of allothetic spatial cues to estimate position based on triangulation [24–26]. These systems continuously reinforce each other, as allothetic cues (especially boundaries and distal landmarks) correct path integration errors and path integration provides a universal metric to construct a framework upon which spatial landmarks can be organized to produce a map [26].

Since spatial locations can be represented by the population activity of place cells, distortion of the mental representation of space might occur if the neural mechanisms that incorporate the allothetic cues onto the map create inhomogeneities in the distribution of place fields. Two-dimensional surface cues often serve as demarcations that segregate the environment into distinct compartments. For example, different tiling on the floor may define the realm of a kitchen and distinguish it from an abutting dining area. Place cells are known to overrepresent apparatus boundaries [27, 28] and goal locations [29, 30], providing evidence of inhomogeneity of the place field map. However, it is not known whether two-dimensional surface cues, which provide no impediment to movement or navigation but which can create a conceptual spatial segmentation of the environment, can also produce inhomogeneity in the map.

To address this question, we trained rats to forage on surfaces with distinct regions demarcated by floor textures or tape line markings. The edges of place fields recorded from dorsal CA1 and CA3 concentrated near the boundaries, resulting in a steeper change in the population vectors of firing rates for locations that cross boundaries compared to locations that do not cross boundaries.

## Results

### Place field edges coincided with the local cue boundaries

We recorded single-unit activity simultaneously from multiple neurons of the CA1 and CA3 pyramidal cell layers of the hippocampus while rats moved clockwise around a circular track in a double rotation, cue-mismatch task (Figure 1A). The quadrants of the track surface were covered by different texture patches (local cues), and objects were placed on the surrounding curtains or on the floor (global cues) [31]. For both standard (STD) and cue-mismatch (MIS) sessions, place fields covered the entire track with no strong tendency to concentrate at specific locations (Figure 1B; Figure S1).

**Figure 1.**
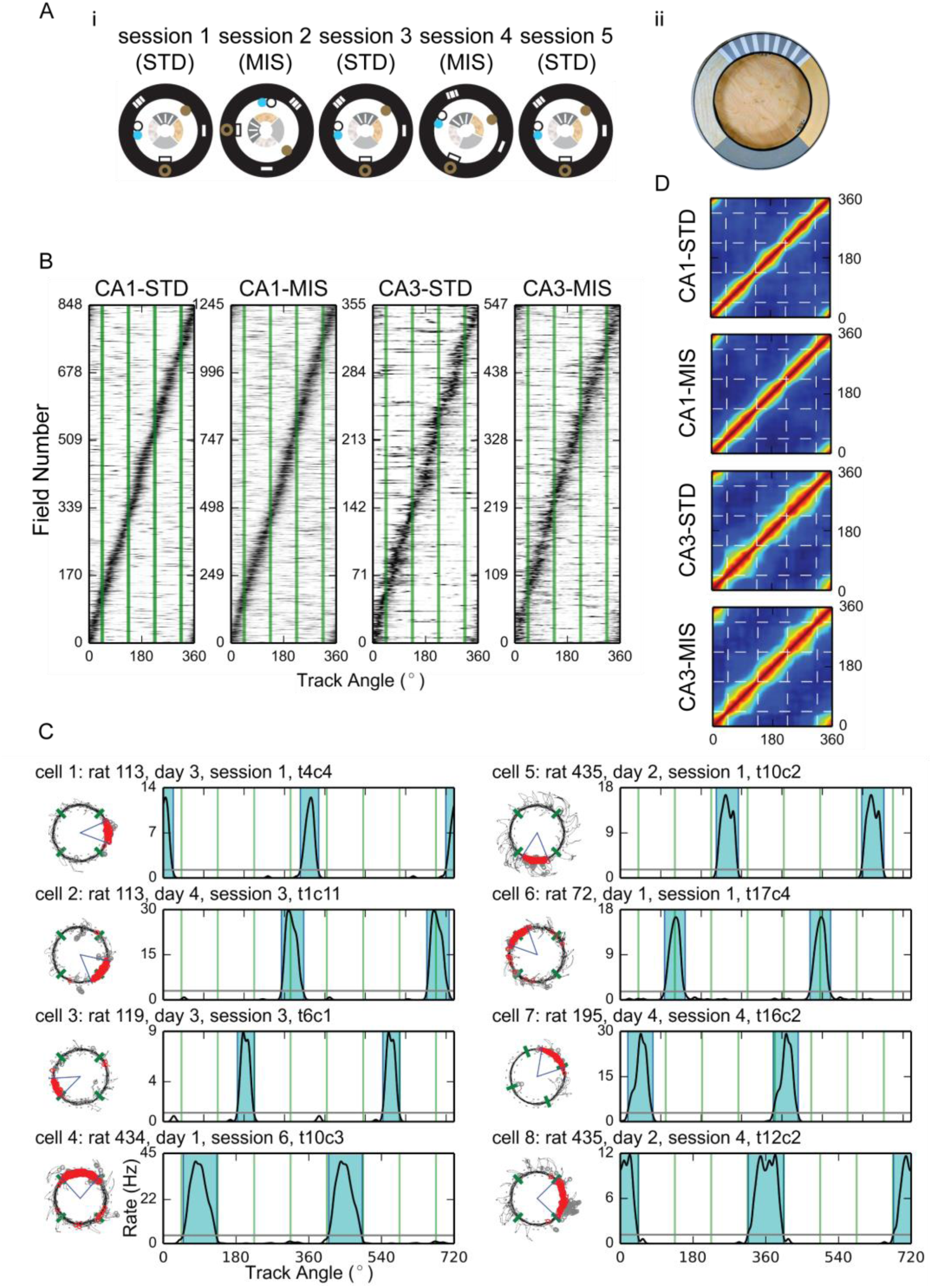
Local-cue boundaries modulated the locations of place field edges. (A) (i) Top-down schematics of the double rotation experiment sessions. Local textures on the circular track are denoted by the different patterns of the inner ring. Global cues are denoted by shapes on the black outer ring representing the black curtains surrounding the track. In this example, 180° (session 2) and 45° (session 4) mismatch (MIS) sessions were interleaved with 3 standard (STD) sessions. (ii) Photograph of the textured, double-rotation track. (B) Sorted firing-rate maps of all the place fields included in the analyses. The abscissa of the map is the track angle and each row of the map is the firing-rate map of a unit. The locations of the local-cue boundaries are denoted by the green lines. The rate maps were normalized by the peak firing rates of each unit and were sorted by the centers of mass of the fields. The same rate map is included in the figure multiple times if the place cell had multiple place fields. (C) Examples of place fields observed in CA1 (cells 1–4) and CA3 (cells 5–8). Some fields confined within a texture quadrant (cells 1 and 5) or crossing a local-cue boundary (cells 2 and 6) had no edges close to any of the local-cue boundaries. However, other fields had one edge near a boundary (cells 3 and 7) or had both of their edges near the boundaries (cells 4 and 8). Fields that had one or more edges near a boundary could be contained within a single texture quadrant or could span across multiple quadrants. For each cell the trajectory-spike plot (left) and the linearized firing-rate map (right) are presented. The blue-shaded areas represent the range of the place field, and all rate maps are duplicated and concatenated in order to show fields crossing 0°. The local-cue boundaries are labeled by green lines in both plots. (D) The cross-correlograms of the PVs. The narrow pinch points of the diagonal band near the local cue-boundaries (dashed lines) show that the PVs changed more rapidly across the boundaries than across similar distances within a texture quadrant. The firing-rate maps were normalized by the peak firing rates before constructing the cross-correlograms; similar results were obtained with nonnormalized matrices (not shown). See also Figure S1 and Figure S7.

**Figure S1.**
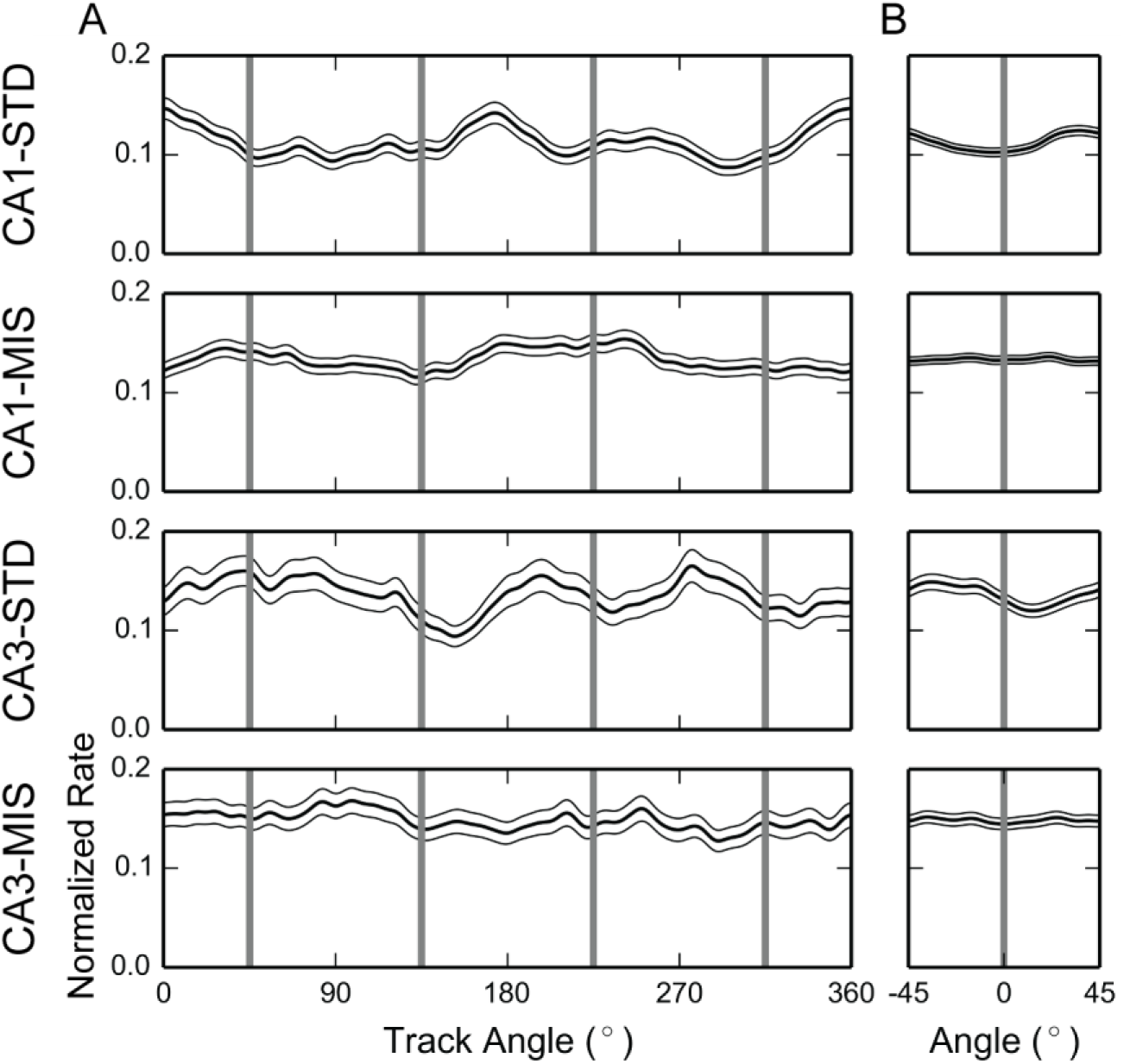
Related to Figure 1; Population firing rates did not change robustly near the local-cue boundaries. (A) The population firing-rate maps. The smoothed firing-rate maps of different units were normalized by the peak firing rate of each unit and stacked together. The mean firing rates across units were calculated and denoted by the thick black curves, and the standard errors were denoted by the thin black curves. The abscissa of the map is the track angle. The gray lines indicate the locations of the local-cue boundaries. There is no clear evidence of changes in mean firing rates at the local-cue boundaries. (B) Local population firing-rate maps near the local-cue boundaries. To further examine whether the population firing rates consistently increased or decreased near the local-cue boundaries, we collapsed the data for each cell across the 4 local-cue boundaries. The behavior and spike data were binned based on their relative locations to the closest boundaries, and the mean firing rates were calculated. The abscissa of the map is the relative location to the local-cue boundary, and the means and the standard errors of firing rates were denoted as in (A). The gray lines label position 0.

Although many place fields crossed local-cue boundaries or fired at a distance from them, there appeared to be a disproportionate number of fields with edges near the local-cue boundaries (Figure 1C). To illustrate this phenomenon, we constructed cross-correlograms of the population vectors (PVs) of firing rates (Figure 1D). The width of the diagonal band reflects the distance the animal must travel before two locations are represented by uncorrelated population activity [32]. The diagonal bands became narrower near the locations of the local-cue boundaries (especially in CA3), indicating a more rapid change in the population activity near the boundaries.

To statistically determine whether more place field edges than expected by chance were located near the local-cue boundaries, we created histograms of the locations of the field edges (Figure 2, left column). The proportions of field edges located ± 15° from the local-cue boundaries were significantly greater than shuffled distributions (Figure 2, middle column, significant for all session types, with significance level α = 0.05, two-tailed test with Bonferroni correction). To provide further support, we performed a bootstrap analysis on the data sample by randomly resampling, with replacement, the same number of place fields that constituted the data set. The field edges of the resampled fields were used to calculate the field edge proportion for each of 1,000 bootstrap trials. Since the local-cue windows occupied 1/3 of the track circumference, we expected to see bootstrapped distributions centered near 0.33 under the null hypothesis of a homogeneous distribution. However, all the bootstrapped field edge proportions were greater than 0.33 (Figure 2, right column, bootstrap confidence intervals with significance level α = 0.05, with Bonferroni correction: CA1-STD, [0.338, 0.396]; CA1-MIS, [0.352, 0.396]; CA3-STD, [0.353, 0.438]; CA3-MIS, [0.371, 0.436]). Both starting and ending edges of place fields revealed a tendency to concentrate near local-cue boundaries (Figure S2A, B), but statistical significance was not reached in all recording conditions (unlike the combined analysis presented in Figure 2). In contrast to the local-cue boundaries, field edges did not appear to concentrate near the global-cue boundaries (Figure S2C). The concentration of place field edges does not appear to be an artifact caused by behavioral biases (Figure S3) or by overrepresentation of place field centers of mass (Figure S4A).

**Figure 2.**
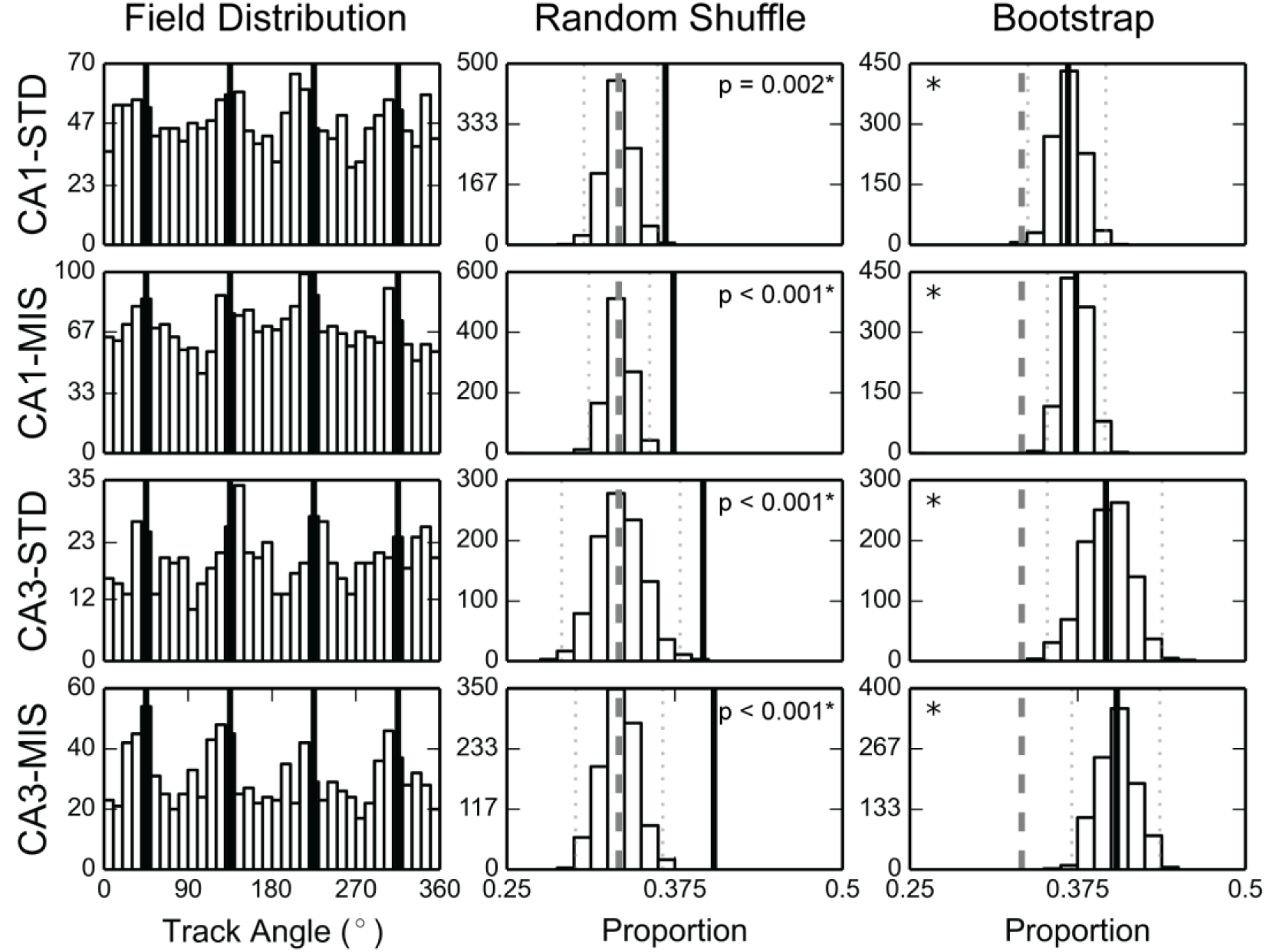
Place field edges coincided with local-cue boundaries. (Left) The distributions of place field edges peaked near the local-cue boundaries (denoted by the black lines). The abscissa of the map is the track angle and the ordinate is the number of field edges observed within the corresponding spatial bin. (Middle, Right) The random shuffling control distributions (middle column) and the bootstrapped distributions (right column) of the proportion of field edges observed within the local-cue windows. The experimentally observed values are denoted by the thick black lines, the 95% confidence intervals of the shuffled distributions by the dotted lines, and the chance level (0.33) by the dashed lines. *, significant at α = 0.05, Bonferroni corrected for 4 comparisons. See also Figure S2, Figure S3 and Figure S4(A).

**Figure S2.**
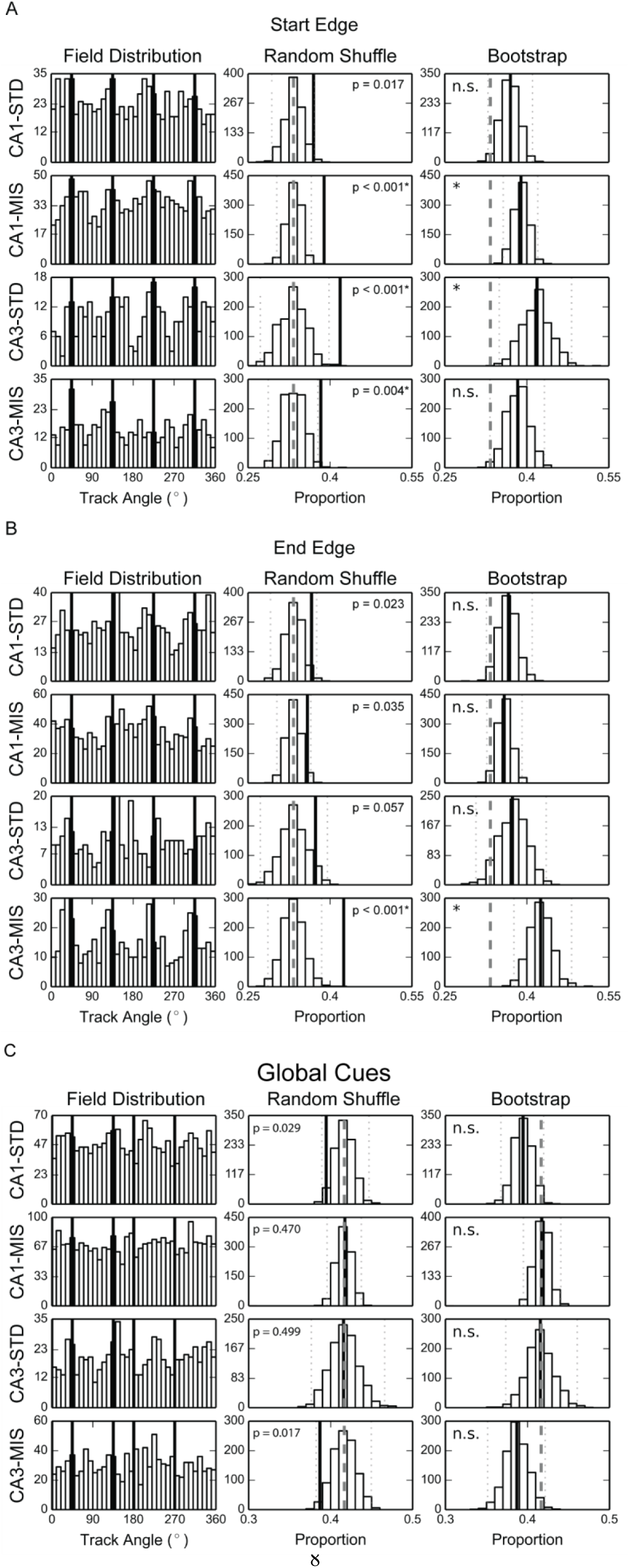
Related to Figure 2; Place fields tended to start and/or end near the local cue boundaries but not the global cue boundaries. (A) (Left) The distributions of the starting edges of the place fields with respect to the local-cue boundaries. (Middle) Shuffling tests were significant at familywise p < 0.05 for all but the CA1-STD session (two-tailed, Bonferroni corrected). (Right) Bootstrap confidence intervals with significance level α = 0.05, Bonferroni corrected: CA1-STD, [0.329, 0.411]; CA1-MIS, [0.357, 0.420]; CA3-STD, [0.350, 0.482]; CA3-MIS, [0.333, 0.432]. Although not all distributions met the stringent Bonferonni-corrected alpha (0.0125 for each distribution), they were all showed the same strong trend. (B) (Left) The distributions of the ending edges of the place fields with respect to the local-cue boundaries. Although the ending edges showed a trend to be preferentially located at the local-cue boundaries, in most cases this tendency did not survive the Bonferonni-corrected statistical tests. (Middle) Shuffling test, significant result for CA3-MIS, with α = 0.05, two-tailed with Bonferroni correction. (Right) bootstrap confidence intervals with significance level α = 0.05 with Bonferroni correction: CA1-STD, [0.326, 0.410]; CA1-MIS, [0.327, 0.391]; CA3-STD, [0.307, 0.435]; CA3-MIS, [0.376, 0.482]. As with the starting edge analysis (A), all distributions showed the same strong trend. (C) (Left) The distributions of the place field edges (start and end) with respect to the global-cue boundaries (i.e., the locations on the track that correspond to the radial angle of the center of each global cue). The solid lines denote the locations of the global-cue boundaries (two local cues were also positioned at 45**°** and 135°). (Middle) Shuffling tests were not significant at familywise α < 0.05, two-tailed with Bonferroni correction. The chance level is ∼0.42 for this analysis since there were 5 global cues and the global cue windows occupied about 42% of the track surface. Note that the observed proportions are almost significantly *less than* the shuffled distributions for the CA1-STD and CA3-MIS sessions; this result is likely explained by the tendency of the peaks to be in the locations of the local cues instead of the global cues. (Right) Bootstrap confidence intervals with familywise α < 0.05 with Bonferroni correction: CA1-STD, [0.368, 0.420]; CA1-MIS, [0.395, 0.441]; CA3-STD, [0.374, 0.461]; CA3-MIS, [0.351, 0.422]. Note that the global cues were large objects distant from the track, and thus the global cue boundaries as defined here were markedly different from the unambiguously defined local cue boundaries. It is thus possible that the global cues had an effect on the place field edges that we were unable to detect in our analyses.

**Figure S3.**
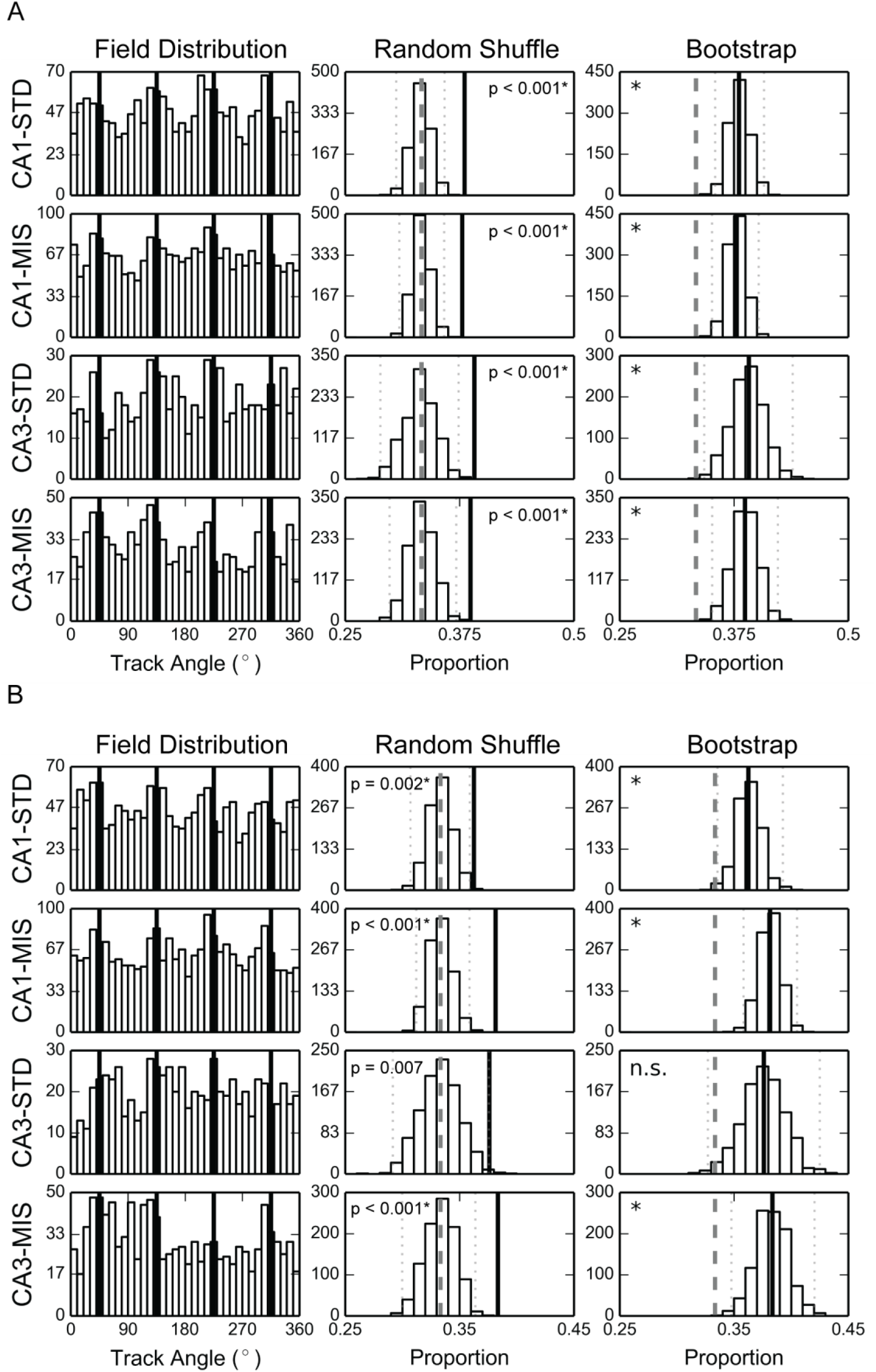
Related to Figure 2; The prevalence of place field edges near local-cue boundaries was not a speed-related artifact. Some rats tended to slow down or pause at the texture edges, a tendency that introduces a potentially confounding variable. It is known that place field firing rates can be modulated by the animal’s momentary running speed, which could affect the precise locations where place field edges were calculated. Two control analyses were performed to address whether inhomogeneities in running speed accounted for the main results on the circular track. (A) The distributions of the place field edges based on the raw data that were not velocity-filtered. Standard practice in the place cell literature is to remove from analysis spikes and position samples that occur when a rat is moving below a threshold speed, in order to discard potential nonspatial firing of cells when a rat is immobile and the hippocampus is in the large irregular activity (LIA) state of EEG. Selective removal of these data points at the local cue boundaries might have artifactually produced the local cue boundary effect on place field edges. To test this, we reanalyzed the data without the speed threshold filtering. For all session types, the shuffling test results and the bootstrap results were still significant with α = 0.05, two-tailed with Bonferroni correction; the bootstrap confidence intervals with significance level α = 0.05, Bonferroni corrected for 4 comparisons: CA1-STD, [0.354, 0.408]; CA1-MIS, [0.351, 0.402]; CA3-STD, [0.342, 0.439]; CA3-MIS, [0.351, 0.423]. (B) The distributions of the place field edges after excluding the data segments in which the rat paused their forward movements. Some rats tended to pause and produce “head scanning” behaviors at the local cue boundaries, a behavior that might have altered the firing of the cells in the place field. To test for this potential confound, we deleted from analysis all traversals across the local cue boundaries in which the rat paused or performed a head scan (see Methods). For all but the CA3-STD sessions, the shuffling and bootstrap results were significant with α = 0.05, two-tailed with Bonferroni correction; the bootstrap confidence intervals: CA1-STD, [0.336, 0.393]; CA1-MIS, [0.358, 0.405]; CA3-STD, [0.327, 0.425]; CA3-MIS, [0.348, 0.420]. These results suggested that the concentration of the field edges near the local-cue boundaries was not an artifact of the tendency of the animal to slow down or pause near the local cue boundaries. The figure formats are as described in Figure 2.

**Figure S4.**
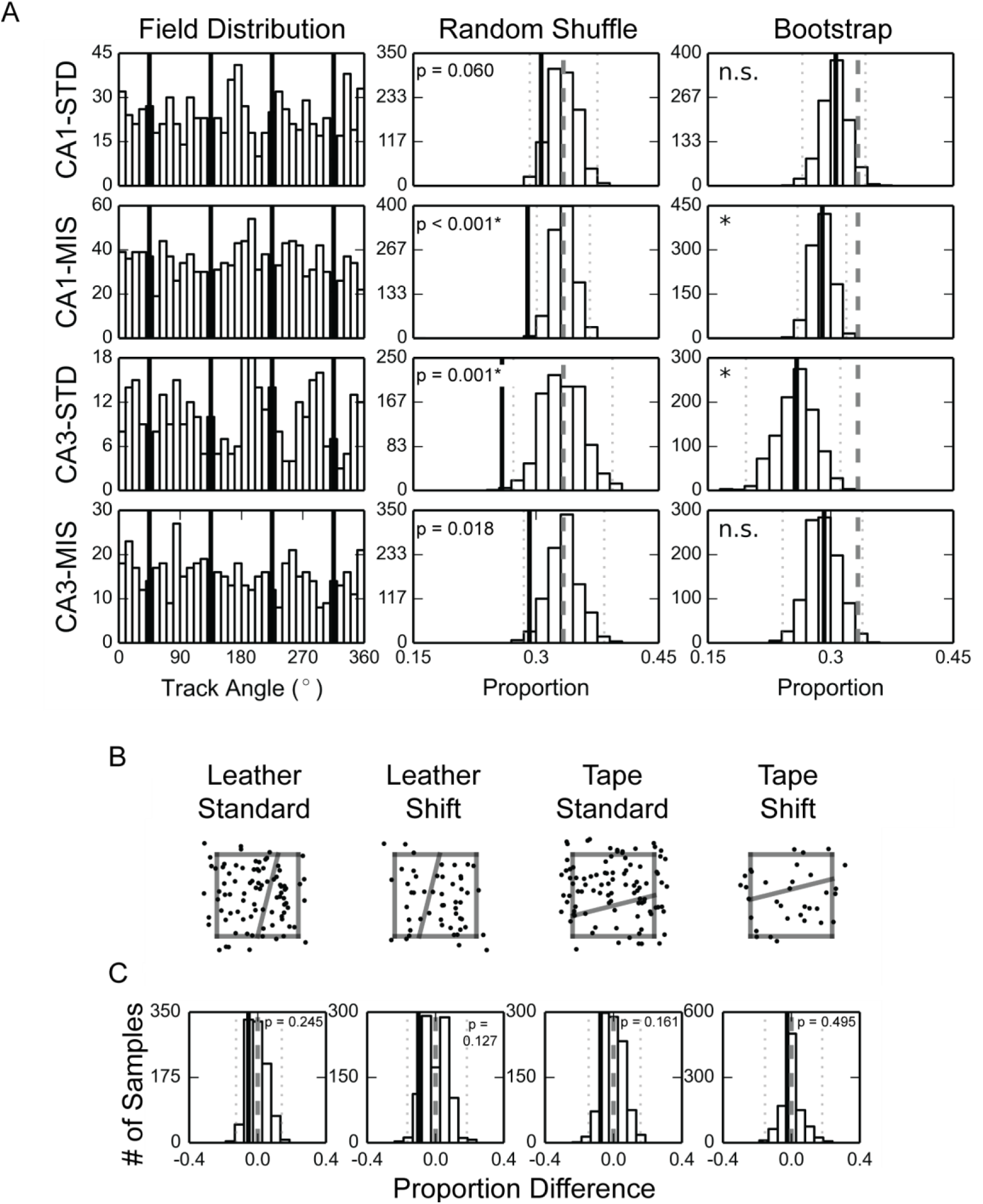
Related to Figure 2 and Figure 5. Surface-cue boundaries were not over-represented by place field centers of mass (CoMs). (A) (Left) The distributions of the place field CoMs on the double cue-rotation track. (Middle, Right) We performed the same shuffling and bootstrap tests as described previously. In contrast to the field edge effects, the observed proportions were significantly fewer than the shuffling results (or trended in that direction) (middle, significant for CA1-MIS and CA3-STD with α = 0.05, two-tailed, with Bonferroni correction) or the uniform distribution chance level (right, bootstrap confidence intervals with significance level α = 0.05, with Bonferroni correction: CA1-STD, [0.265, 0.343]; CA1-MIS, [0.259, 0.319]; CA3-STD, [0.197, 0.312]; CA3-MIS, [0.241, 0.339]). This underrepresentation of field COMs near the local-cue boundaries might be explained by the sizes of place fields. Given that 70% of the CA1 and CA3 place fields were 33°-103° in length and the local-cue boundaries were at 90° intervals, most fields that started or ended near the local-cue boundaries would have their COMs away from the local-cue boundaries. The greater-than-chance prevalence of place fields with edges near the local-cue boundaries would therefore lead to a low prevalence of COMs near the boundaries. (B) The distributions of place field CoMs on the simple boards. Each dot represents the CoM of a field. The proportions of fields that had CoMs within the boundary zones on the simple boards were not significantly different from the plain-board control (two-tailed χ^2^ test with d.f.=1: leather-standard, χ^2^= 0.900, p=0.343 (n.s.); leather-shift, χ^2^= 2.118, p=0.146 (n.s.); tape-standard, χ^2^= 1.500, p=0.221 (n.s.); tape-shift, χ^2^= 0.144, p=0.704 (n.s.). α=0.05 with Bonferroni correction for 4 comparisons.) (C) A permutation test suggests the cue boundaries were not over-represented by place field CoMs. The proportions of place fields with their CoMs located within the boundary zones were calculated, and a permutation test as performed in Figure 5B was used to compare the field proportion differences between the simple boards and the plain board control. In all cases the data fell within the 95% confidence intervals of the permutation distributions. The denotations are as described in Figure 2.

### Firing rate maps modulated by cue boundaries in 2-dimensional environments

Place field properties can be different when rats run stereotyped trajectories on one-dimensional (1-D) circular or linear tracks, compared to when they perform more irregular foraging in two-dimensional (2-D) open fields or platforms [33]. We thus examined whether place field edges concentrated near surface texture boundaries in 2-D environments. We first trained 6 rats to forage on a *complex board* with a complicated surface pattern composed of geometric shapes constructed from different texture patches and tape lines (Figure 3A). CA1 place cell recordings from 5 rats show that some place field edges were aligned with a subset of the cue boundaries and corners (Figure 3B). These examples provide compelling visual demonstrations that the place field edges respect the local texture boundaries on the platform, similar to the circular track. A number of place fields crossed some boundaries, even as they were aligned to other boundaries; thus, as on the circular track, the fields were not always contained within a single bounded region. However, the complexity and heterogeneity of place field edges relative to the complex geometric patterns on the board precluded a detailed quantitative analysis.

**Figure 3.**
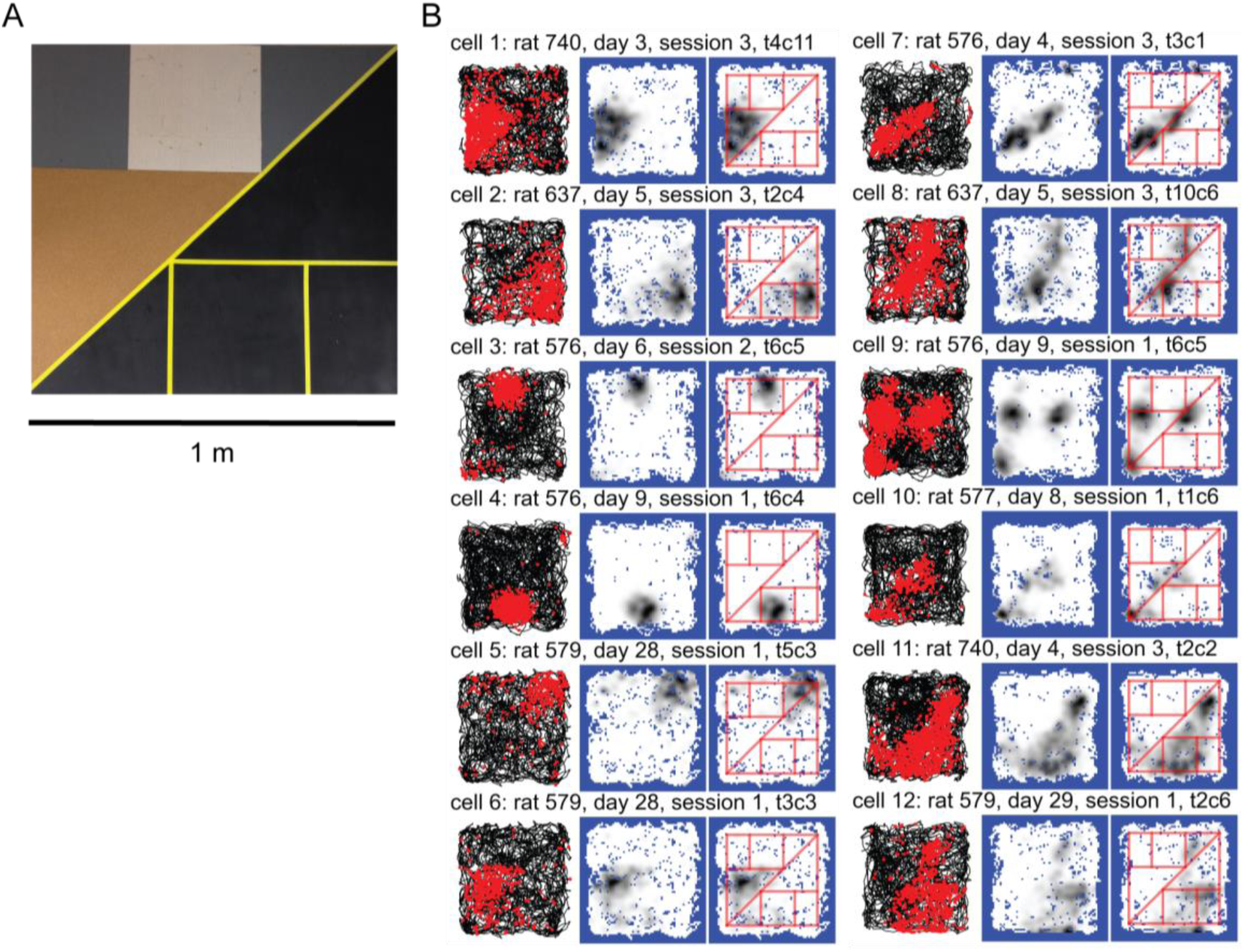
Place field edges modulated by surface boundaries on the complex board. (A) Photo of the complex board. (B) Examples of CA1 place fields modulated by the surface boundaries. For each cell the trajectory-spike plot (left), the smoothed firing-rate map (middle), and the smoothed firing-rate map with superimposed cue boundaries (right) are presented. The cue boundaries appeared to modulate the edges of the place fields: Cells 1-6 occupied one or multiple geometric shapes defined by the cue boundaries, and they developed triangular, rectangular, or complex-shaped fields. Cells 7-8 fired along one or more cue boundaries and had elongated, stripe-like fields. The cue boundaries also appeared to affect the number and locations of the place fields. For example, cell 9 developed 3 fields at the vertices of the brown triangle and cell 10 fired at the corresponding corners of the black triangles. Cells 11 and 12 developed complicated firing patterns. The field of cell 11 filled in the black area, with some “bleeding” into the lower-left of the textured area, and the field of cell 12 occupied the rectangular area near the bottom but also extended along the diagonal boundary.

To quantify place field alignment to 2-D boundaries, we collected further data from 3 rats foraging on *simple boards* with a single cue boundary. The simple board experiments contained two types of boards. The *leather boards* contained a cue boundary formed by the contrast between a leather surface patch and a wooden texture; the *tape boards* contained a white tape line that divided the board into two sections (Figure 4A). The surface patterns of the leather and the tape boards were 180°-rotated, mirror images of each other. Therefore, any possible field edge concentration effect observed for the latter board could not easily be explained by the effect generated for the preceding board. The experiments consisted of 2 consecutive sessions with the texture boundary in a standard location, a shift session in which the boundary was moved to a new location, and a final standard session. Place fields were distributed over the entire surface of the simple boards and a subset of the fields appeared to be modulated by the boundaries (Figure 4B).

**Figure 4.**
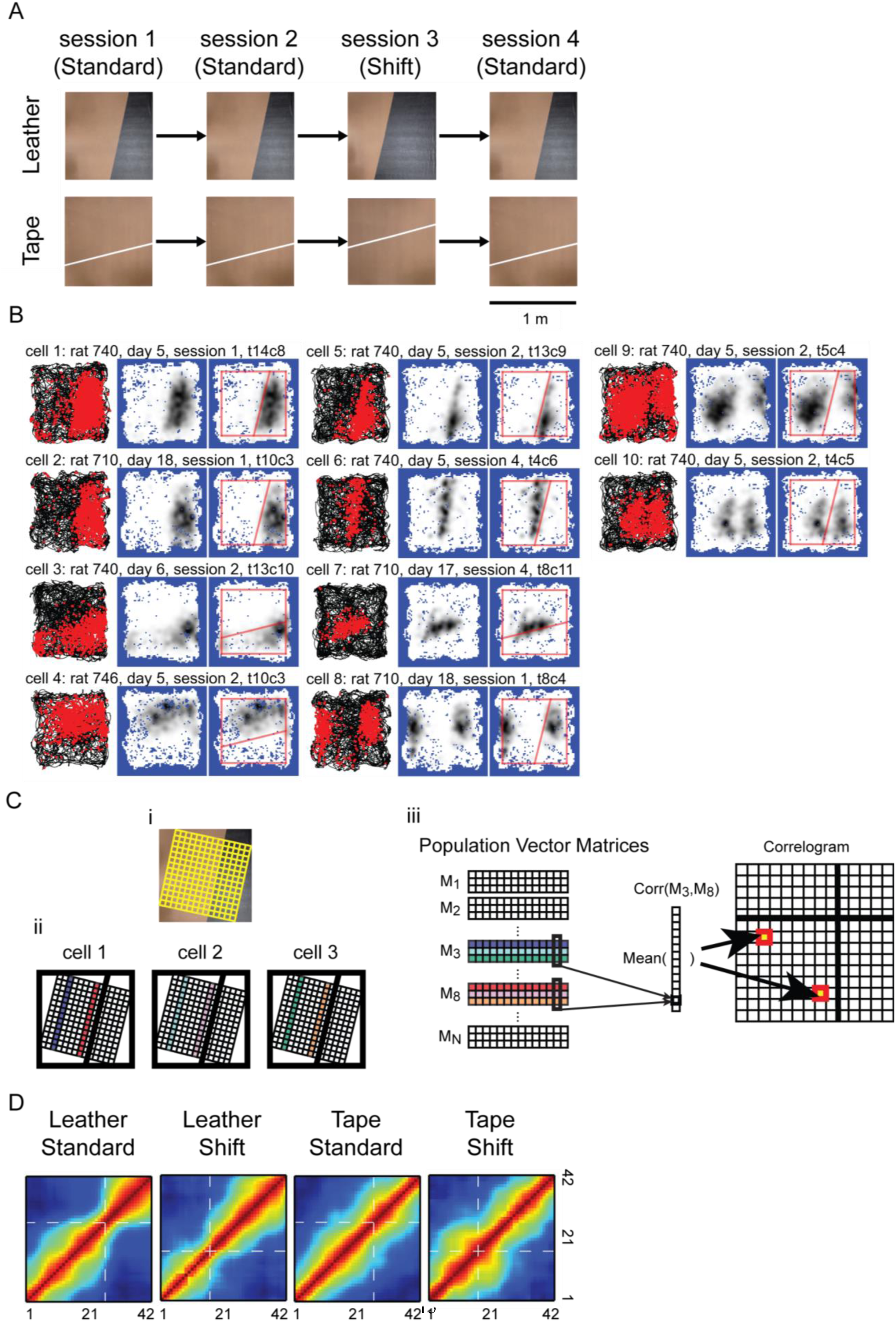
Place field edges modulated by surface boundaries on the simple boards. (A) Photos of the simple boards and schematics of the simple board foraging task protocol. (B) Examples of CA1 place fields modulated by the cue boundaries. The figure format is as described in Figure 3. Similar to the complex board, we observed place fields that occupied geometric shapes defined by the cue boundary and the edges of the experiment board (cells 1-4), as well as elongated fields extending along the cue boundary (cells 5-7). Some of the neural correlates near the cue boundary were similar to those near walls or other traditionally defined boundaries reported in previous studies. Cell 8-9 resembled boundary cells in that they developed multiple fields at corresponding locations with respect to the cue boundary and the board edge. However, for cell 9 the two fields were different in size, which might reflect the reset of cell activity at the cue boundary, or imply that the cell has one large field that was split by the boundary. Similarly, cell 10 seemed to have two fields that were intersected by the boundary, as if its firing activity was inhibited by the cue boundary. Only a small number of cells had firing patterns similar to cells 8-10. (C) Schematics of the cross-correlogram construction process for the simple boards. (i) The spatial binning of the simple board. The firing-rate map was constructed based on binning (denoted by the yellow grid) aligned with the surface boundary. (ii)(iii) The construction of the cross-correlogram. Assuming there were only three cells (denoted by the squares in (ii)), the firing rates of the third columns of the grids from each cell (the purple, cyan, and green stripes in (ii)) were stacked together and formed the third PVM (M3 in (iii)), and the 8^th^ columns (the red, pink, and orange stripes in (ii)) formed M8, etc. For each element of the cross-correlogram, the Pearson’s product-moment correlation coefficients were calculated between the corresponding columns of the selected PVMs, and the mean correlation was calculated across the columns. (D) The cross-correlograms of the PVs. Along the direction perpendicular to the surface boundary, the correlation dropped more abruptly near the surface boundaries (denoted by the white dashed lines) for the leather boards. The firing-rate maps were normalized by the peak firing rates before constructing the cross-correlograms. See also Figure S5 and Figure S7.

To visualize whether the field edges were modulated by the cue boundaries at the population level, we partitioned the simple board into 42 equally-spaced stripes parallel to the cue boundary and calculated the similarities between the population activity vectors of the stripes (Figure 4C). The widths of the diagonal band (warm colors) of the normalized cross-correlograms decreased near the cue boundary locations for the leather-standard board (similar to the narrowing at the texture boundaries of the circular track in Figure 1B), but they remained relatively homogeneous along the diagonal lines for the tape-standard board (Figure 4D). Similar results were obtained for the nonnormalized correlograms, although some inhomogeneity of the diagonal band width started to appear for the tape-standard board (data not shown). When the cue boundary changed location, many place fields remapped between the standard and shift sessions (Figure S5). Nonetheless, the diagonal bands were narrowed at the cue boundaries for the leather-shift board. These results indicated that the place cell population activity for the leather boards changed more abruptly for two locations across the cue boundaries than for equivalent distances within a texture, while the effect was much weaker for the tape boards.

**Figure S5.**
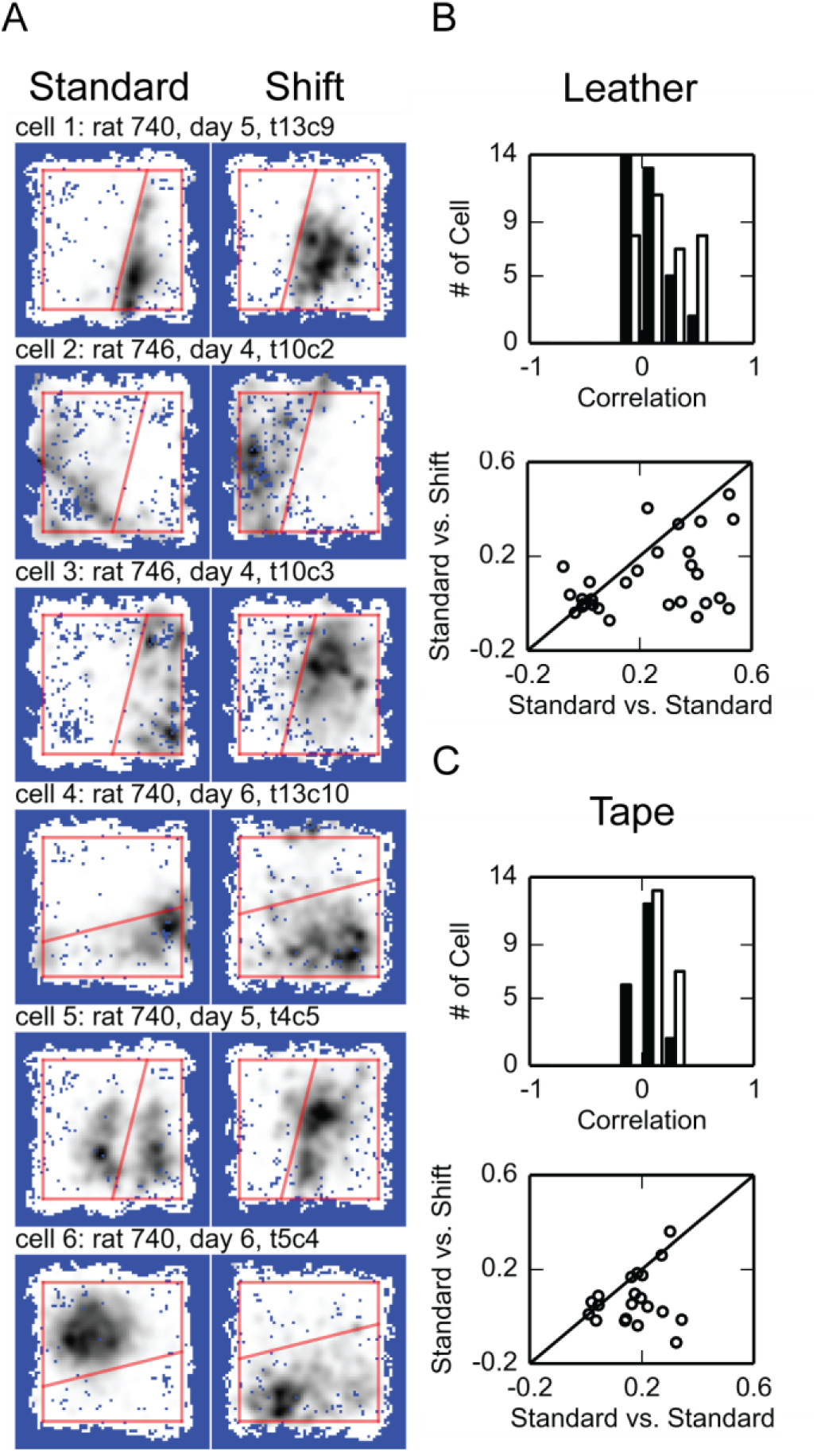
Related to Figure 4; Shift of cue boundary triggered place cell remapping. (A) Examples of corresponding place field changes when the cue boundary was shifted to a new location. Each row represents the data collected from the same cell in the shift session (right) and the preceding standard session (left). Upon the manipulation, there were fields with their edges following the boundary (cells 1-4), fields separated by the cue boundary merging into one field (cell 5), or fields flipping along the cue boundary (cell 6). (B) Remapping was observed between the standard and the shift session for the leather board. (Top) the distributions of the correlations between firing-rate maps from different sessions. The standard vs. standard correlation coefficients are denoted by the white bars, and the standard vs. shift correlation coefficients by the black bars. The median of the standard vs. standard correlation coefficients was significantly larger than standard vs. shift (n = 34; two-tailed Wilcoxon signed-rank test: T = 102.000, p = 0.001; with significance level α = 0.05, with Bonferroni correction for 2 comparisons). (Bottom) The scatter plots of the correlation coefficient pairs. The abscissa of the scatter plot is the correlation coefficient between two standard sessions and the ordinate is the correlation coefficient between the standard and the shift session. Most points were below the diagonal line, which indicated that for most units the correlation coefficients were larger when comparing between the standard sessions than comparing between the standard and the shift session. (C) Remapping was observed between the standard and the shift session for the tape board. The figure formats are as described in (B). The correlation coefficients were larger when comparing between the standard sessions than comparing between the standard and the shift session (n = 20; two-tailed Wilcoxon signed-rank test: T = 33.000, p = 0.008; with significance level α = 0.05, with Bonferroni correction for 2 comparisons).

### Place field edges concentrated near the boundaries on the simple boards

To determine statistically whether place field edges concentrated at the leather or tape boundaries, similar to the texture-cue boundaries on the circular track (Figure 2), we created 2-D field-edge density maps for the 4 types of boards. Hot spots along the cue boundaries were observed for all the boards (although much weaker on the tape-shift board) (Figure 5A), demonstrating a trend for the field edges to concentrate near the cue boundaries (as well as near the perimeters of the boards). We calculated boundary preference indices (BPIs) to quantify whether high field-edge densities were more frequently observed in the “boundary zone” (i.e. locations that were ≤ 10 cm from the cue boundary), than in the “nonboundary zone” (i.e. locations that were > 10 cm from both the cue boundary and from the periphery of the board).

**Figure 5.**
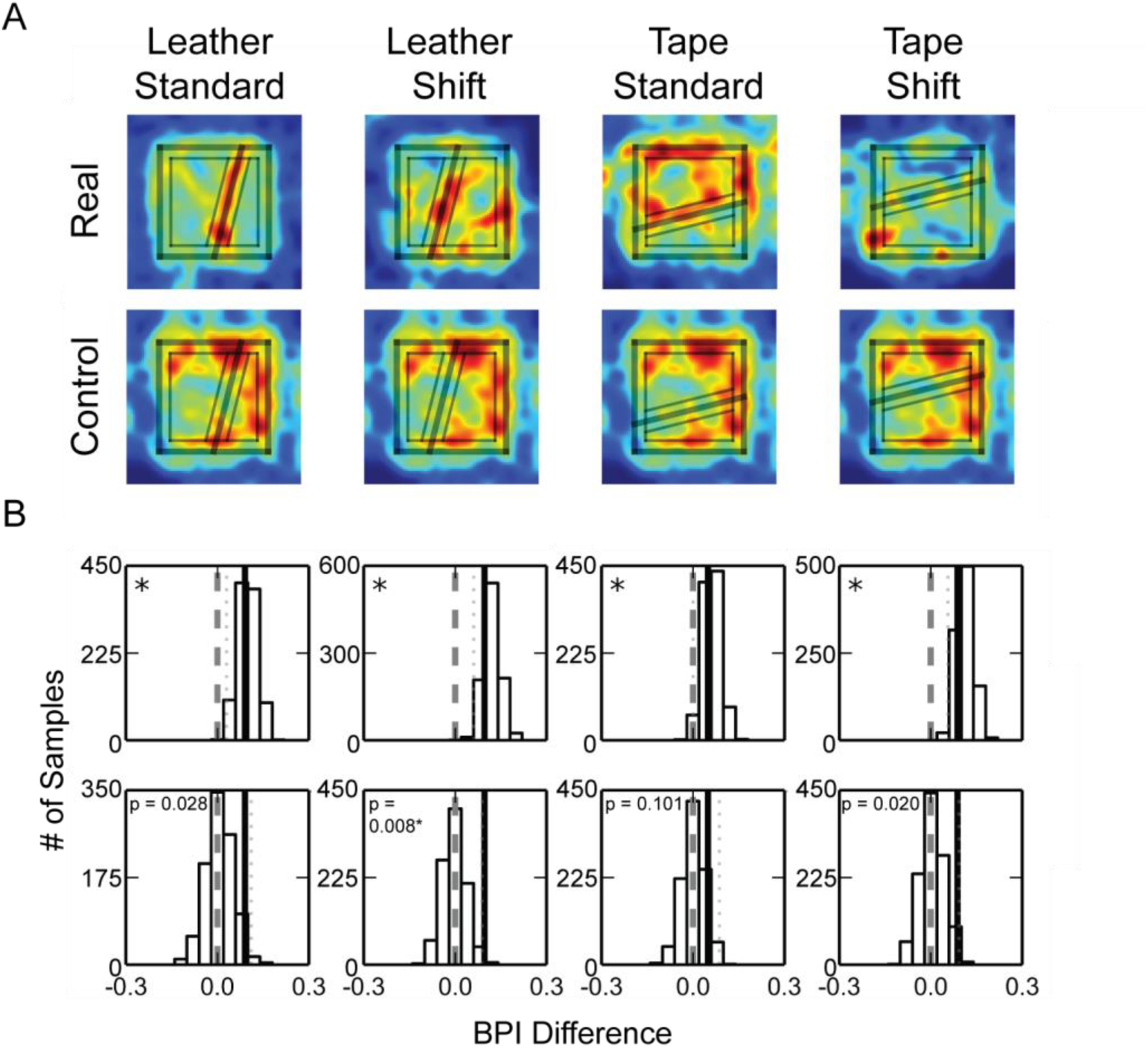
Place field edges concentrated near the surface boundaries. (A) The place field edge density maps of the simple boards (top) and the control plain board with zone markings from the corresponding simple boards (bottom). The rim of the platform and the cue boundary are labeled by thick lines, and the boundary and non-boundary zones used in the analyses are labeled by thin lines. (B) The bootstrapped distributions (top) and the permutation distributions (bottom) of the BPI differences. The figure formats are as described in Figure 2. See also Figure S4(B).

For each board we randomly resampled the place field identifiers with replacement to create 1000 bootstrapped samples of place field subsets, from which a BPI was calculated for each place field subset. Concurrently, we projected the boundary and nonboundary zone partitions of the simple board onto a control board with a plain surface and calculated the BPI of the data collected from the *plain board* accordingly. By randomly pairing observed and control bootstrapped samples, we calculated the BPI difference between the selected samples (leather/tape board – plain board control).

The BPI differences were significantly larger than zero for all boards (Figure 5B; a one-tailed test was used because we had a strong, *a priori* prediction based on the results of the circular track experiment; one-tailed cut-off value of the bootstrapped distribution with significance level α = 0.05, with Bonferroni correction: leather-standard, 0.030; leather-shift, 0.061; tape-standard, 0.000; tape-shift, 0.056). This result implied that, for all conditions, the field edges were more concentrated near the cue boundaries than could be expected by chance. A permutation test showed similar trends, although only the leather-shift board attained statistical significance (significance level α = 0.05, one-tailed, with Bonferroni correction).

### Adjacent place fields extended along the cue boundaries

The field edge concentration effect can be a result of (a) a disproportionate number of fields neighboring the cue boundary, (b) elongated field edges along the cue boundary, or (c) a combination of these possibilities (Figure 6A). For all simple boards, the proportions of place fields that overlapped completely or partially with the boundary zone were not different from the plain board (Figure 6B), and the cue boundaries were not overrepresented by the place field centers of mass (Figure S4B, C). On the other hand, the field edge lengths within the boundary zones, defined as the number of spatial bins within the boundary zones containing the edge of a specific field, were on average significantly greater than the plain-board control for the leather boards (Figure 6C, top row). The edge length analysis excluded place fields that did not overlap with the boundary zones. Similar results were obtained with a permutation test (Figure 6C, bottom row). Thus, for the standard boards, we observed higher field-edge density differences than expected by chance because the place fields close to the cue boundaries tended to extend along the boundaries, thereby increasing the length of the field edge aligned with the cue boundaries.

**Figure 6.**
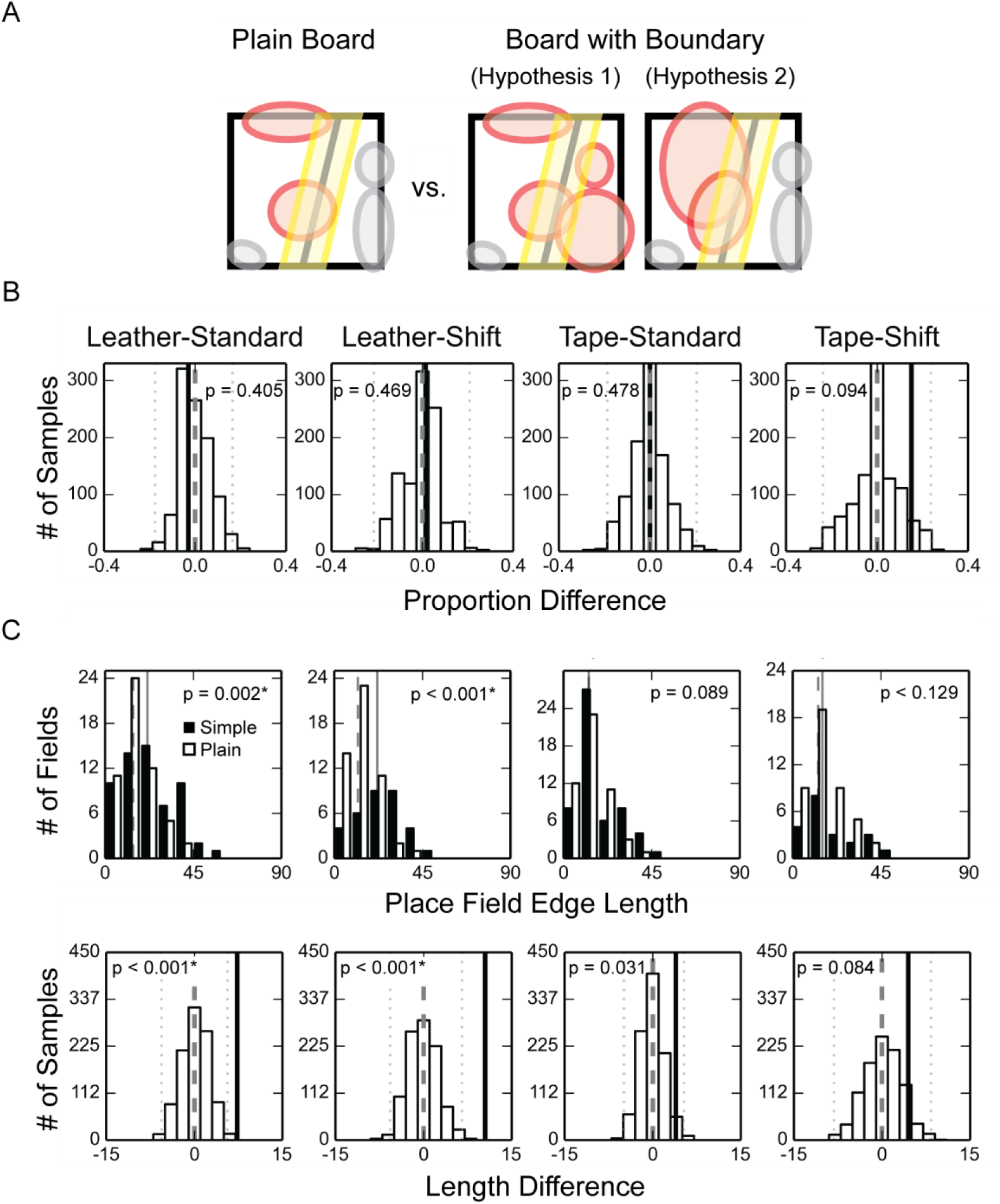
Adjacent fields extending along the surface boundaries. (A) Schematics of different hypotheses explaining why the BPI difference was larger on the experiment boards than on the plain board. Hypothesis 1 suggests that a higher proportion of place fields were observed within the boundary zone on the simple boards than on the plain board. Hypothesis 2 suggests that place fields tended to extend along the cue boundary on the simple boards, and thus the average length of field edges observed within the boundary zone was larger on the simple boards than on the plain board. The place fields are denoted by colored circles, and the red fields increase the field edge density within the boundary zone. (B) The permutation test of the field proportion difference (no boards pass significance test at α = 0.05, two-tailed, with Bonferroni correction). The denotations are as described in Figure 2. (C) Longer field edges were found near the surface boundaries on the leather boards than on the plain board. (Top) The distributions of the field edge lengths within the boundary zone. The distributions of the field edge lengths collected from the simple boards are represented by black bars, with the median values denoted by the solid lines; and the distributions of the plain board control are represented by white bars, with the median values denoted by the dash lines. The field edge length of the leather boards were significantly larger than the plain board control (two-tailed Mann-Whitney U test, n is the number of fields, and m is the median of the contour lengths: leather-standard, ntexture = 59, nplain = 54, mtexture = 21.53, mplain = 14.27, U = 1099.0, p = 0.002*; leather-shift, ntexture = 33, nplain = 51, mtexture = 21.93, mplain = 12.06, U = 397.0, p < 0.001*; tape-standard, ntexture = 54, nplain = 50, mtexture = 12.94, mplain = 12.92, U = 1142.5, p =0.089; tape-shift, ntexture = 21, nplain = 44, mtexture = 15.22, mplain = 13.07, U=381.0, p = 0.129; α = 0.05 with Bonferroni correction). (Bottom) The permutation test of the edge length difference. The denotations are as described in Figure 2.

**Figure S6.**
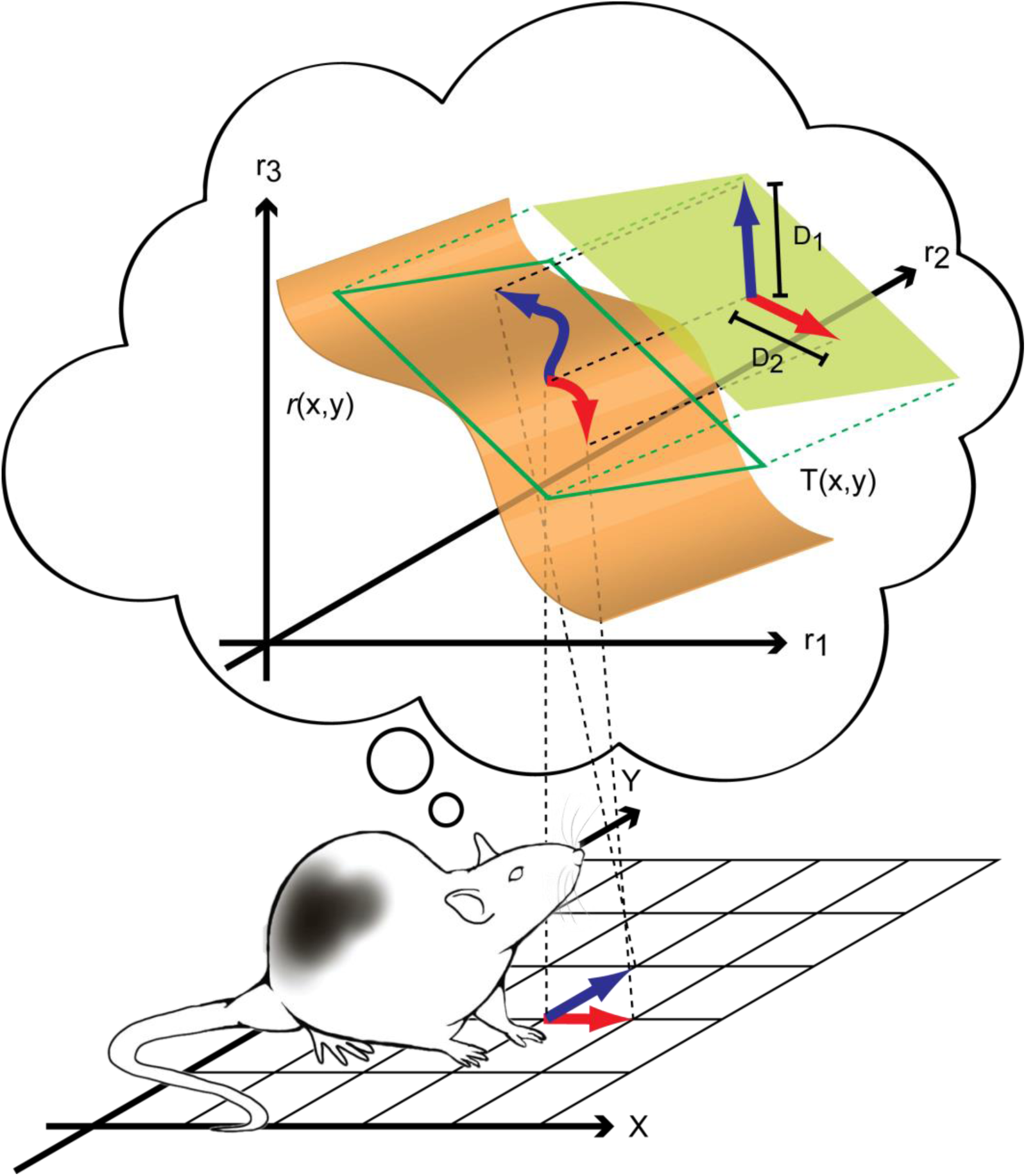
Related to Figure 7; Interpolation of PV. A linear approximation method was used to interpolate the PV differences. On the experiment platform (represented by the x-y plane at the bottom), the PV can be empirically calculated for any spatial bin. The PVs are discrete samplings of the continuous function r(x, y) (the orange surface) which defines the PVs at arbitrary locations. However, these Cartesian bins limit the calculation of PV differences to 8 directions (i.e., the difference between a bin and the 8 surrounding bins). In order to examine the differences between PVs in directions more fine-scaled than the Cartesian bins allow, for every spatial bin, we computed the tangent plane T(x, y) (the green plane) of r(x, y) and interpolated the PV for arbitrary locations. The red and blue angles denote two orthogonal movement directions in the physical world, and their corresponding PV changes in the neural space. D1 and D2 are the resulting PV differences (defined as the Euclidean distances) for these two different movement directions. See Methods for more details.

**Figure 7.**
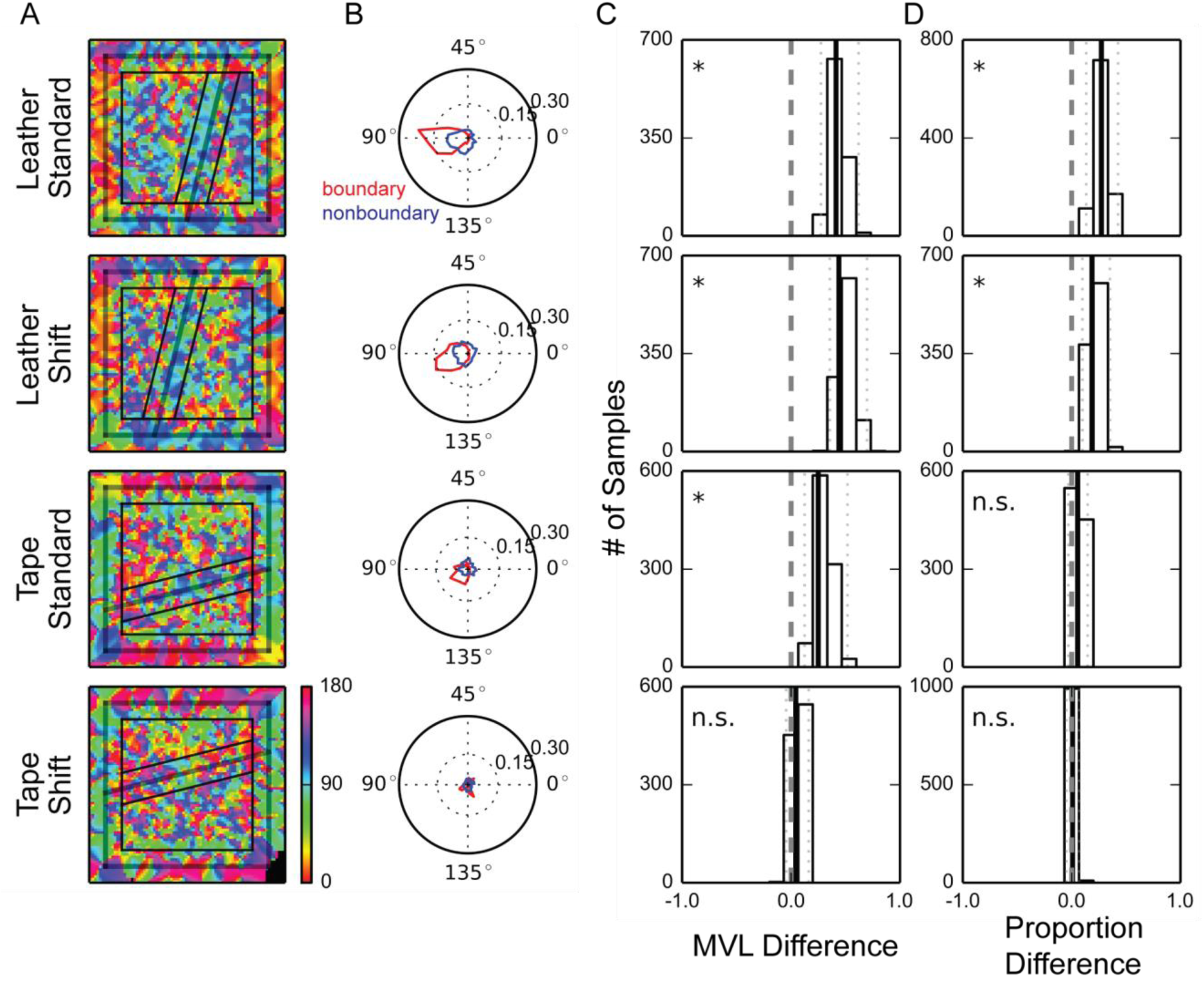
Population activity changes more abruptly across, not along, the surface boundary. (A) Visualizations of the directions along which the PVs changed most abruptly. Red represents the direction parallel to the surface boundary and blue represents the direction perpendicular to the boundary. The denotations of the lines are as described in Figure 2. (B) The Rayleigh plot of the directions observed within the boundary zone (red) and the non-boundary zone (blue). The movement direction ranged from 0° to 180°, 0° when parallel with the cue boundary and 90° when perpendicular to the cue boundary. (C) The bootstrap distributions of the MVL differences. (D) The bootstrap distributions of the spatial bin proportion difference. The denotations of (C) and (D) are as described in Figure 2. See also Figure S6 and Figure S7.

### Population activity changes more abruptly across, not along, the boundary

The structures of the place field edge distributions described in the previous section suggested that the PVs at adjacent locations would be more similar when the locations were both within a texture area than when the locations were across a texture boundary. To examine this hypothesis, for each location we calculated the direction along which the PVs changed most (Figure 7A; Figure S6; see Methods). For all but the tape-shift board, the changes in PVs were the largest in the direction perpendicular (or near perpendicular) to the cue boundaries for most locations within the boundary zone; for the nonboundary zone, the directions were more divergent (Figure 7B). To test this difference statistically, we first examined whether the directions of maximal PV difference were more concentrated in the boundary zone than in the nonboundary zone. We computed the difference between the mean vector lengths (MVL) of the direction distributions of the two zones. When the rate maps were normalized by the peak firing rates before the construction of the PVs, bootstrapped distributions were significantly larger than zero for all but the tape-shift board (Figure 7C, bootstrap confidence intervals with significance level α = 0.05, with Bonferroni correction: leather-standard, [0.275, 0.619]; leather-shift, [0.357, 0.697]; tape-standard, [0.125, 0.518]; tape-shift, [-0.045, 0.161]). Similar results were obtained when the PVs were constructed based on the raw firing-rate maps (data not shown).

To determine whether the direction bias was preferentially perpendicular to the direction of the cue boundary, we calculated the proportion of spatial bins in which the directions were ±15° from the angle perpendicular to the cue boundary. Bootstrapped distributions of the difference in this proportion between the boundary and nonboundary zones were significantly larger than zero for the leather boards; a similar trend was observed for the tape-standard board but was not significant (Figure 7D, bootstrap confidence intervals with significance level α = 0.05, with Bonferroni correction: leather-standard, [0.134, 0.426]; leather-shift, [0.099, 0.352]; tape-standard, [-0.030, 0.147]; tape-shift, [-0.047, 0.070]). When the PVs were constructed based on the raw firing-rate maps, the bootstrapped distribution was significantly larger than 0 for the tape-standard board as well (data not shown). These results suggest that for the leather boards (and weakly for the tape-standard board), the directions of maximal population decorrelation were significantly more consistent and perpendicular to the cue boundaries when the animals were near the boundaries than when the animals were away from the boundaries.

## Discussion

In the current study, we showed that the edges of place fields tended to concentrate near internal surface boundaries when rats foraged on an apparatus covered by inhomogeneous surface patterns. The field edge concentration phenomenon was observed on both a 1-D circular track and 2-D open platforms, and it could be elicited either by boundaries between different surface textures or (weakly) by a tape line. These results demonstrate that rats not only use surface texture cues as reference points to anchor the orientation of the cognitive map, as shown in previous studies [31,34–36], but they also encode the locations of the surface texture cue boundaries. The tendency for individual place cells to switch on or off near the cue boundaries sharpened the differences between the population vectors (PVs) of firing rates across a cue boundary and differentiated the representations on either side of the boundary.

### Representation of spatial segmentation in the cognitive map

The present study can be contrasted with prior studies of hippocampal correlates of spatial segmentation that have investigated how place cells distinguish similar, connected environments [37–43]. In these studies, a significant number of place cells repeated their firing patterns in geometrically corresponding locations across perceptually similar compartments with high walls oriented in the same direction. Analogous repeating firing patterns were also seen in grid cell maps when rats ran through a hairpin maze [44] or on early exposure to an environment consisting of two visually identical boxes connected by an external corridor [45]. In these experiments, the compartments were perceptually and geometrically similar and repetitive. Therefore, it is perhaps not surprising that similar sensory inputs within each compartment would trigger the same units of the cognitive map to fire at corresponding locations [46–48]. Although path integration could, in principle, have provided overriding input to distinguish the compartments, this influence appears to have fostered differences in firing rates of the place fields across compartments rather than creating completely new representations. However, when the compartments were oriented differently, place cells were able to distinguish the compartments and did not repeat their firing fields, presumably because the head direction cell system was able to discriminate the orientation of the boundaries across the compartments [39].

The experiments in the present paper are similar to these prior experiments in that we investigated how the hippocampus represents geometrically segregated segments of a larger space. Our experiments differ, however, in that the spatial segments were defined not by high-walled boundaries but by changes in the texture of the surface upon which the rat moved. Unlike the high-walled environments, unique views of the global environment were attained from different textured segments. In this case, only a few cells had multiple place fields that appeared to be located in geometrically similar subareas (Figure 4, cell 8-9). Nonetheless, spatial segmentation could still be deciphered by examining the locations of the place field edges.

### The “geometric module”: Dissociation between spatial segmentation and reorientation

Another series of studies relevant to the current work is the investigation of the “geometric module” in rodents and humans. When asked to find hidden rewards in a high-walled rectangular environment, human children tend to search at both the correct and the geometrically equivalent locations even when these two locations can be distinguished by non-geometric features [4950] (but see [51]). These results suggest that human children used the geometry of a space to solve the task. Similar results have been observed in rodents (in which the phenomenon was first described) [52–54], birds [55, 56], and fish [57]. When the walls were replaced by small curbs, the boundaries (i.e. curbs) no longer defined the perceptually and navigationally available environment. Nonetheless, human children and birds still used the geometry of the segment to solve the task [50–56], suggesting that the spatial segment was recognized as an isolated part of the environment and possessed geometric features.

However, not all internal boundaries provide geometric information for reorientation purposes. When the environments were segregated by a luminance contrast on the floor (e.g., a black rectangle painted on a white floor), human children and chickens no longer made systematic errors at the geometrically equivalent location [50–56], and imaging of the parahippocampal place area and retrosplenial cortex showed weaker responses compared to boundaries that extended into the z axis [58]. One might conclude that these flat surface boundaries were undetected by the cognitive mapping system, thereby precluding the influence of the geometric module. Our results demonstrated, however, that even when the spatial compartments were segregated by flat surface cues, the demarcation information was present in the cognitive map (although perhaps inaccessible to the spatial orientation system). Thus, information about the *presence* of geometric boundaries may be dissociated from the *use* of this geometric information for orientation. Although the influence of environmental geometry on head direction cell tuning is influenced by complex factors [59–61], it is nonetheless possible that the geometry-controlled reorientation phenomenon is caused in large part by geometric control over these cells. In turn, the head direction cells can reorient downstream grid cells and place cells by virtue of their close coupling [62–64]. Alternatively, the reorientation may be largely dependent on boundary-selective neurons [16-18,65], which may not respond to the floor texture boundaries of the present study (although Figure 4 shows two examples of cells that are similar to a boundary/border cell). In any case, the reported inability of organisms to reorient to geometric boundaries defined by flat surfaces does not necessarily imply that the spatial representations of these shapes are not encoded.

### Concentration of field edges may elongate the mental distance across a boundary

These results may help explain certain phenomena from the human literature regarding distorted representations of space. When requested to memorize the locations of objects or landmarks in a compartmented environment, people tend to underestimate distances between targets within the same spatial compartment and overestimate distances between targets in different compartments [4,6–9,66]. Similarly, judgment errors in relative spatial relationships increase when two objects are located in different spatial compartments [5, 67]. Even when there are not explicit boundaries (i.e., physical barriers, markings, or discrete transitions in context) in an environment, subjects tend to exaggerate the distance between two locations if the locations are in two conceptually distinct regions connected by a smooth transition (e.g. a woods gradually changing into a field) [4, 6]. The systematic errors in estimating angles or distance suggest that the psychological representation of space is not isomorphic to physical space, and our results might provide insight into the underlying physiological mechanisms instantiating local distortions of the mental representation.

There are a number of well-studied neural coding schemes that the hippocampus might have used to represent the surface texture boundaries in our experiments. First, the hippocampus might have overrepresented the boundary by developing a larger number of place fields along the boundary, similar to the overrepresentation previously shown for the peripheral borders of an environment [27, 28], starting sites [68, 69], or goal locations [29, 30]. However, our analyses of place field locations provided little evidence for a disproportionate number of place fields located at the texture boundaries. Second, the place cells might have fired at a higher or lower mean rate at the boundaries than in the middle of the textures. Again, our analyses showed little evidence of such rate coding of the boundaries. Instead, the surface texture boundaries appeared to be encoded at the population level by a more abrupt decorrelation of the ensemble representation of space as the rat crossed the boundary, compared to when it moved an equivalent distance in the center of a texture segment or along a boundary. A downstream structure able to decode the rate of change of the neural representation would thus be able to detect the presence of the boundary.

At a local scale, the magnitude of correlation between the neural representations of different locations reflects the physical distance between the locations. In an environment where the place fields are homogeneously distributed, if two locations are not farther than the average size of place fields (i.e., the spatial scale factor), the distance between them would be negatively correlated with the similarities between their neural representations [70]. However, if the distribution of place fields is inhomogeneous, such that the correlation between PVs of neighboring locations can vary, the mental distance between these locations might vary accordingly. In our data, the surface cue boundaries were encoded by a concentration of place field edges, and this representation decreased the correlation between the PVs across the boundaries. We therefore hypothesize that a ‘mental gap’ would be inserted in the animal’s perception of distance traveled whenever an animal moved across or mentally traversed through the boundary (Figure S7).

The insertion of mental gaps may also elongate the perceived distance between locations at a more global scale. When two locations are sufficiently far apart, such that there is no longer any overlap in the population of place cells encoding the locations, the representations are maximally decorrelated with no further relationship to longer distances. However, the brain may estimate distance between two remote locations by integrating distances between neighboring points connecting these locations. The mental distance between any two locations across the surface texture boundary may thus be elongated, since they would be connected by paths including the mental gap. It has been shown in other sensory systems that a local change near the boundary can elicit a global perceptual effect. For example, the Cornsweet Illusion [71] demonstrates that when two areas of equal brightness are separated by two local illumination gradients at the border, the entire areas are perceived as having different brightnesses defined by the strong contrast that exists only at the border. The perception of spatial segregation might similarly be mediated by neural mechanisms that can extend the mental gaps generated at the boundaries to regions farther from the border, creating a global percept of greater distance across the entire environment.

**Figure S7.**
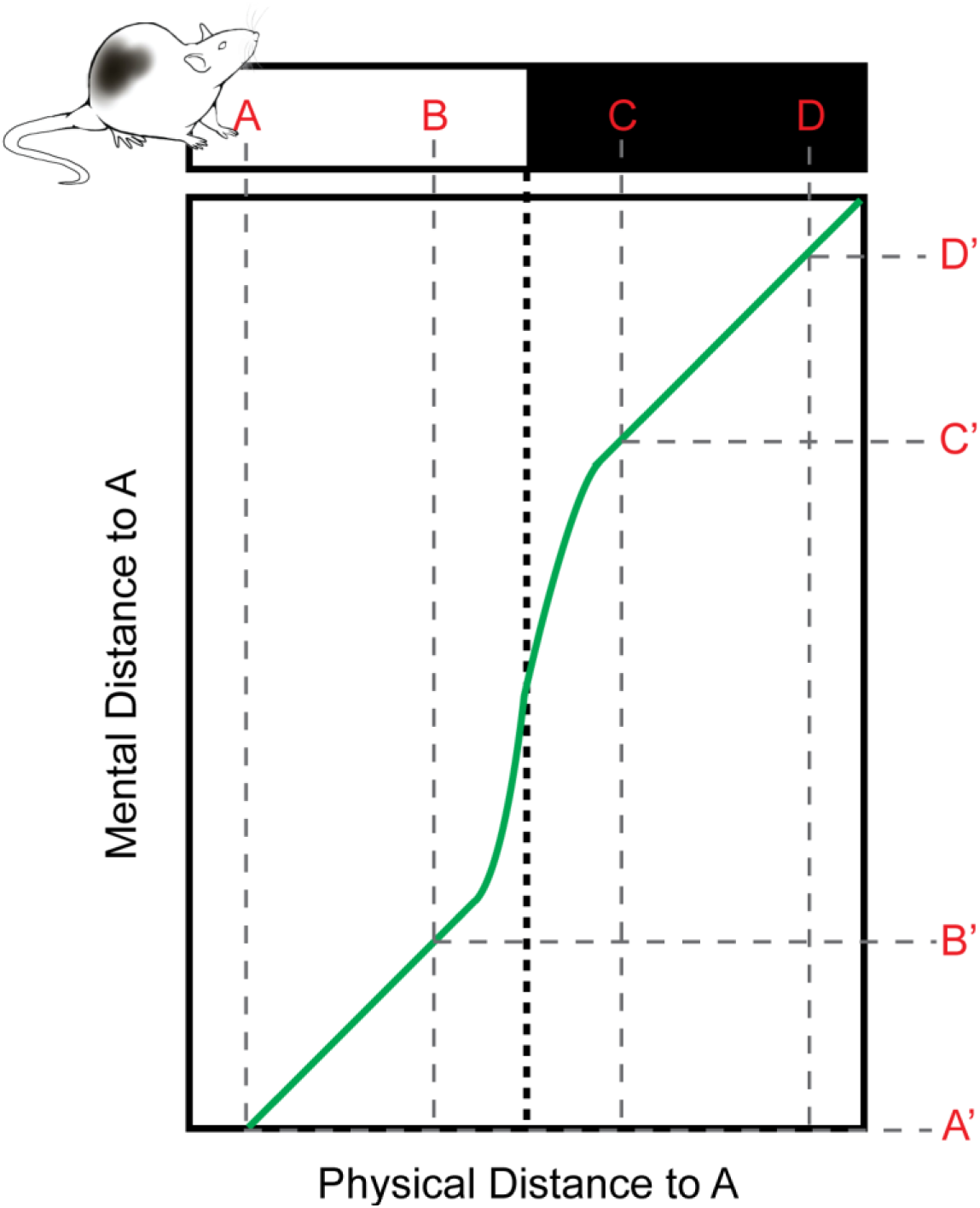
Related to Figure 1 and Figure 4; A global perceptual effect can be triggered by a local boundary. A schematic of the hypothesized relationship between the actual distance traveled by a rat and the corresponding perceived distance based on the mental gap hypothesis. On a linear track (denoted by the rectangle on the top) covered by two different surface textures (represented by the black and the white areas), a mental gap is hypothesized to be inserted when the rat crosses the texture boundary, and the perceived travel distance would thus be longer than its physical length. Although the exaggeration of perceived distance occurs only locally near the boundary (between point B and C), the distortion of distance perception can be a global effect (perceived distance is also elongated between point A and D)

## Methods

### Subjects and surgery

A total of 49 adult male Long-Evans rats were used in this study: 41 rats participated in the double rotation task, 6 rats in the complex-board forage task, and 3 rats in the simple-board forage task (see below for task descriptions). Separate groups of rats were used in different tasks except for one rat that underwent both the complex-board forage task and the simple-board forage task. The double rotation data were previously collected and published for other purposes [72–79]. The rats were housed individually on a 12/12-h light/dark cycle and all experiments took place during the dark phase of the cycle. The rats had free access to water but were food restricted such that their body weights were maintained at 80-90% of the *ad libitum* level.

For surgical implantation of a microdrive array, the rat was injected with ketamine (60 mg/kg) and xylazine (8 mg/kg), followed by isoflurane inhalation to produce a surgical level of anesthesia. A craniotomy was made on the right hemisphere, and the microdrive array was placed at the center of the craniotomy targeting the dorsal hippocampus. For post-operative analgesia, the rat was administered ketoprofen (5 mg/kg) or meloxicam (1 mg/kg) subcutaneously, or 1 cc of oral acetaminophen (Children’s Tylenol liquid suspension, 160 mg) right after the surgery. Further analgesia was either provided on the following two days by meloxicam administered orally or blended in food (Metacam, 1∼2 mg/kg), or provided on the following day by a second injection of ketoprofen or by access to diluted acetaminophen in drinking water as needed. All implanted rats received 0.15 ml of enrofloxacin (Baytril, 2.27%) and 30 mg of tetracycline blended in food daily until termination. All animal procedures complied with U.S. National Institutes of Health guidelines and were approved by the Institutional Animal Care and Use Committee at Johns Hopkins University or the University of Texas Health Science Center at Houston.

### Electrophysiology and recording electronics

Microdrive arrays that contained 6-20 independently adjustable tetrodes were built for extracellular recordings. Each tetrode was composed of four 12 or 17 μm nichrome wires, or four 17 μm platinum-iridium wires, twisted together. The tips of the nichrome wires were individually gold-plated to reach 200-500 kΩ impedance measured at 1 kHz. After at least four days of recovery from surgery, each tetrode was advanced gradually per day over 20-40 days until its tip arrived at the hippocampal CA1 or CA3 layers and activities of pyramidal cells were observed while the rat rested on a pedestal.

During recording, the neural signal was buffered by a unity-gain preamplifier and filtered between 600 Hz and 6 kHz by the data acquisition system (Neuralynx, Bozeman, MT). Whenever the electrophysiological signal passed a threshold between 50-70 µV, a 1 ms segment was extracted at 32 kHz and stored as a spike waveform. To track position, the head stage was equipped with protruding arms extending backwards or to the sides of its head. Red and green light emitting diodes (LEDs) were attached to the arms to track head position and direction, captured at 30-60 Hz by cameras mounted on the ceiling.

### Single-unit isolation

Single units were isolated offline with customized spike-sorting software (Winclust, J. Knierim). For each tetrode, the putative spikes were displayed as points in a multidimensional waveform parameter space, and the points were manually clustered primarily based on the relative spike amplitudes and energy simultaneously recorded from individual wires of the tetrode. The isolation quality was subjectively rated on a scale of 1 (very good) to 5 (poor), representing the extent to which a spike cluster could be separated from other clusters and noise. The ratings were completely independent of any spatial or behavioral correlates of the unit. Units categorized as 4 (marginal) or 5 (poor) were excluded from analyses.

To ensure that we did not artificially inflate the sample size by repetitively sampling the same units across multiple sessions, for each tetrode we only included the day with the largest number of place fields recorded. When the same types of sessions were presented in the same day, we only included the cell-session with the highest within-field mean firing rate of the day in our analyses.

### Histology

After the experiments were complete, the rats were anesthetized with 1 cc of Euthasol and were transcardially perfused with saline followed by 4% formalin. In some rats, a subset of tetrodes was selected to pass current and create marker lesions 24 h before perfusion. After perfusion, the cranium was partially opened and the brain was exposed to formalin for at least 4 h with the tetrodes in place to preserve the tracks of the tetrodes. The brain was extracted and soaked in formalin for 12 h before transfer to a 30% sucrose formalin solution (wt/vol). After the brain was frozen, it was sectioned at 40 μm in the coronal plane and stained with 0.1% cresyl violet. Recording locations of the tetrodes were assigned by matching the identified tetrode tracks on the brain slices against the known configurations of the microarrays and marker lesions, if any. For the tetrodes targeting the CA1 and CA3 layers on different recording days, depth reconstruction of the tetrode tracks was performed for each recording session to identify the brain region from which the units were recorded.

### Double rotation task

#### Protocol

Rats were trained to run clockwise on a circular track (76 cm O.D., 10 cm wide) to collect food pellet rewards placed at arbitrary locations on the track. On average, the rats obtained ∼2 rewards/lap, but this varied across rats and sessions as needed to promote good performance. The recording sessions started after the rats learned to continuously run on the track with few pauses (∼1-2 weeks of pretraining). Before the recording session, the rat was disoriented (by being placed in a covered box and walked a number of cycles around the apparatus) and placed at an arbitrary starting location on the track. The same food reward schedule was used as in the training sessions and the session ended after the rat finished ∼15 laps around the track. In both training and recording stages, whenever the rat turned around and moved counterclockwise, the experimenter would block its path with a piece of cardboard until it turned back and resumed the clockwise movement. The experimenter also discouraged grooming behavior by snapping fingers or activating a hand-held clicker when the rat paused to groom.

The quadrants of the circular track were covered by differently textured surfaces which served as local cues, starting from 12 o’clock and in the clockwise direction: gray duct tape with white tape stripes, brown medium-grit sandpaper, a gray rubber mat with a pebbled surface, and beige carpet pad material [31]. The track was placed in a circular, curtained environment (2.7-m diameter) in which six distinct objects were present either on the floor or on the curtain as global cues. For the standard (STD) sessions, the local and global cue configuration was maintained as during training. For the mismatch (MIS) sessions, the global and local cues were rotated clockwise and counterclockwise, respectively, to achieve total cue mismatches of 45°, 90°, 135° or 180°. Each day of recording consisted of either 5 sessions, with three STD sessions interleaved with two MIS sessions, or 6 sessions, identical with the 5-session day except for an additional STD session at the start. The mismatch angle for each MIS session was pseudo-randomly selected such that each angle was experienced once during the first 2 days and once again during the second 2 days. For most of the rats there were four days of recording, but for a small proportion of rats there were over 10 recording days. We used only the first four days of recording of each rat to balance the data.

#### Spatial cell filtering

A linear classifier based on the average firing rate and spike waveform width was applied to units with isolation qualities in category 1 to 3 to select and exclude the putative interneurons, which have narrower waveforms and higher mean firing rates than principal cells. Units that were identified as interneurons by the experimenter during spike sorting were also excluded. The remaining cells were classified as putative pyramidal cells, and they were included in quantitative analyses if they fired at least 30 spikes during forward movement. For all analyses (unless noted otherwise), data were discarded when the rat was not running forwards (i.e. when its speed was less than 10°/s, when its head protruded beyond the track edge, and when a lateral head-scanning movement or pausing behavior was detected [78]), in order to prevent contamination of the results by nonspatial firing that occurs during immobility.

The standard Skaggs spatial information measure [80] tended to produce false negative errors when applied to 1-D data [78], and thus we also incorporated the Olypher spatial information score [81] to compensate for the Skaggs measure [78]. The statistical significance of both Skaggs and Olypher measures were computed by temporally shifting the spike trains to construct the control distribution. Since the rats were trained to run continuously on a circular track, the temporal sequences of the rat positions were quasiperiodic, and thus we additionally reversed the spike trains before the time-shifting procedure to break the regularity and prevent creating false negative results [78]. For the putative pyramidal cells with enough spikes, cells were analyzed if the Skaggs score was larger than 1.0 bits per spike or the Olypher score was larger than 0.4 bits, and the score was > 99% of the scores from the shuffled data.

#### Place field detection

Place fields were visualized by creating trajectory-spike plots and firing rate maps. The trajectory-spike plot shows the trajectories of the animal (denoted by black curves), and the locations of the animal when a spike was detected (denoted by circles). The running spikes were denoted by red circles while the spikes excluded by the velocity filter were denoted by gray circles. The average firing rates were calculated as the spike counts divided by the occupancy durations within each track-angle bin (1°), and the firing rate vectors were circularly smoothed with a Gaussian kernel with standard deviation 4.3°. Putative place fields were isolated by thresholding the smoothed firing-rate vectors at 10% of the unit’s maximum firing rate and grouping the contiguous bins with firing rates larger than the threshold. Putative fields separated by only 1 track-angle bin were merged. After merging, the fields that had maximum firing rate > 1.5 Hz and that were ≤330° long were included in the following analyses. The large upper bound was chosen based on the observation that a small number of putative pyramidal cells fired almost all over the track but they were silent within a small gap. The median place field size was 60° with interquartile range (IQR) 38° for CA1 fields, and was 73° with IQR 62° for CA3 fields; only two fields were larger than 270° (Fig 1 (b)). The starting and ending edge location of the place fields were defined as the starting and ending track angle bins, respectively.

#### Cross-correlograms

We constructed population firing rate cross-correlograms to visualize the similarities between the place cell population activity recorded at different track locations. Place cells were identified and their firing-rate vectors were calculated and smoothed as described in the 1-D place field construction section. Vertically stacking the transposed firing-rate vectors formed an N x 360 population firing rate matrix, where N is the number of units used to construct the matrix. The i^th^ row of the matrix was the firing-rate vector of the i^th^ unit, and the j^th^ column of the matrix was the population vector (PV) of firing rates at the j^th^ track-angle bin. The cross-correlogram was constructed by calculating the Pearson correlation coefficients between pairs of PVs. The (i^th^,j^th^) element of the cross-correlogram was the correlation between the i^th^ and j^th^ column of the population firing rate matrix [73, 82]. To balance the activity strengths across different units, we reported the normalized cross-correlograms in which the firing-rate vectors of individual units were divided by their maximum firing rates before being stacked together.

#### Place field edge distribution

To examine whether the place field edges concentrated near the local-cue boundaries, we defined 30° wide zones centered on the local cue boundaries as the local-cue windows and calculated the proportions of place field edges located within the windows. Both field shuffling and bootstrap techniques were used to test whether the proportions were significantly higher than chance level.

For the shuffling test, the place field locations were randomly rotated while the field sizes remained the same. For each field the rotation angle was randomly selected from [0°, 360°) and the in-window field edge proportion was calculated based on the rotated field edge locations across the population. This shuffling procedure was performed 1000 times. The distributions of the shuffled in-window field edge proportions simulate the expected distributions assuming the fields were randomly scattered on the track. The result was significant if the percentage of the shuffled samples that were larger than or equal to the observed in-window field edge proportion was smaller than 0.00625 (significance level α = 0.05, two tailed with Bonferroni correction for 4 comparisons).

To bootstrap the data, we randomly resampled with replacement the same number of place fields as the original set 1000 times. In each trial the bootstrapped in-window field edge proportion was calculated to construct the bootstrap distributions. The observed in-window field edge proportions were then compared against the confidence intervals of the bootstrap distributions.

#### Control for head-scanning and pausing behavior

To verify that the field edge concentration effect was not a result of head-scanning or pausing behavior interfering with the place field detection algorithm, we ran a separate control analysis (Figure S3) that excluded a larger range of data when a scan or pause was detected (see [78]) near or within a place field. For each place field, a window which was 15° wider than the field on both sides was defined. If a scan or pause started within the window, we removed the behavior and spiking data of that traversal through the window. After the same removal process was performed on every field, we excluded the cell-sessions if, for any part of the track, no data were left after the deletion. We detected place fields and constructed the field edge distributions based on the filtered data following the same procedures described in the previous sections.

### Complex and simple board tasks

#### Complex board protocol

Five rats were trained to search for chocolate pellets placed at arbitrary locations on an open field with a homogeneous surface texture. The recording experiments started after the rats learned to continuously run on the platform with few pauses. In each recording session, either a textured board with a complex surface pattern (the *complex board*) or a plain board with a uniform surface texture (the *plain board*) was placed at the center of a circular, curtained environment with no deliberate salient global cues. The rats performed the same foraging task for 20 min on the board. For three of the rats, 1-3 *complex board* sessions were performed each day, followed by one *plain board* session in some cases. The other two rats experienced two *plain board* sessions followed by one complex board session per day. There was a minimum of two days of recording for each rat.

The *complex board* was 1 x 1 m and its surface was composed of a complex combination of geometric shapes demarcated by different texture patches and tapes (Figure 3A). The upper left half of the platform was covered by different surface textures: a grey rubber mat with a pebbled surface shaped as a rectangle and a small triangle, white sandpaper shaped as a square and brown cork mat shaped as a large triangle. The lower right half of the platform was uniformly painted black with yellow tape labeling borders 180° rotationally symmetric to the upper left half. The *complex board* was novel to the rats on the first day of recording. The *plain board* was a 1.1 x 1.1 m wooden board and its surface was uniformly painted black.

#### Simple board protocol

The same training procedure as describd in the *complex board protocol* section was used to train three rats to forage in an open field. For each 20 min recording session, the rats performed the same foraging task on a textured board with a slanted linear boundary crossing the surface. The board was placed at the center of a circular, curtained environment with no deliberate salient global cues.

Four different boards were used in the *simple board forage task*: *leather-standard*, *leather-shift*, *tape-standard*, and *tape-shift*. The brown smooth wooden surface of each leather board was partially covered by a black, synthetic leather patch, and the boundary between the two surface textures was an oblique line crossing the board. For the leather-standard board, the separation line passed the bottom edge of the board at the center, and the top edge at 25 cm from the top-right corner. For the leather-shift board, the separation line shifted 20 cm to the left. The brown wooden surface of each tape board was labeled by an oblique white tape line crossing the board. The geometric patterns of the tape boards were 90° rotated mirror images of the leather boards. Furthermore, while wooden surfaces were present on both the leather and tape boards, the textures of the surfaces were different, in that the leather board had a smoother wooden texture than the tape board (Figure 4A). These features reduced the possibility that a place cell would fire at the same location across different boards.

There were two days of recording for each rat, and the rats foraged on the leather boards for one day and on the tape boards for the other day. For two out of three rats the leather boards came first. During each recording day, the rat experienced two *standard* sessions, followed by a *shift* session and back to the standard session. The rat was brought out of the experiment room for 5-10 min between sessions to rest and was provided access to water on a pedestal. All four boards were novel to the rats before the first recording session of the board, and for two rats the simple boards were the first experiment apparatus with inhomogeneous surfaces (the other rat performed the complex board foraging task before the simple board foraging task). The data collected from the *plain board* (See *Complex board protocol*) were used as the control data for the simple board forage task.

#### Firing-rate map construction

The experiment boards were divided into small spatial bins and the average firing rate at each bin was smoothed to construct the firing rate map. We binned the experiment boards in different ways as described in the corresponding sections depending on the purposes of the analyses. The average firing rates were calculated as the spike counts divided by the occupancy duration within each spatial bin. Only running activities with velocity > 5.76 cm/sec (to match the velocity filter used in our double rotation task) were included in spatial cell analyses.

A Gaussian kernel with standard deviation 3 cm was applied to the average firing rates, and the smoothed firing-rate maps were then used in the quantitative analyses. Similar results were obtained when an edge preserving smoothing algorithm, which adopted both Kuwahara [83] and median [84] smoothing filters, was used (results not shown).

#### Spatial cell filtering

The isolated units were scrutinized as described above to exclude putative interneurons. The spike trains of the units were considered reliable only if the isolation quality was at category 3 or better, and there were at least 50 running spikes recorded in the session. The Skaggs spatial information scores were calculated for the qualified cell-sessions to examine whether their firing activities were spatially tuned. For each cell-session, an area extending 30 cm beyond each side of the experiment board (to capture firing when the rat’s head was off the board) was partitioned into a matrix of 2 x 2 cm spatial bins. The smoothed average firing rate for each bin was calculated as described in the *Firing-rate map construction* section, and the spatial information and the p value were calculated based on the smoothed firing-rate maps. To pass the spatial-cell criteria, the cell-session must have spatial information ≥ 0.6, at a significance level of 0.01.

#### Cross-correlograms

In order to compare the population neural activities of the place cells across the cue boundaries, we binned the simple boards with grids that were aligned with the cue boundaries (not orthogonal to the board edges) (Figure 4Ci). To maximize available data without including the out-of-platform area, a rotated square area inscribed in the platform rim was used for the analysis. The rotated square was ∼ 82.5 x 82.5 cm^2^ and was divided into a 42 x 42 matrix with each bin ∼ 4 cm^2^.

For each cell-session, the averaged firing rates of the spatial bins were calculated as the spike counts divided by the occupancy durations, and the firing-rate maps were smoothed as described in the *Firing-rate maps construction* section. Each column of the firing-rate map corresponded to a band parallel to the cue boundaries (Figure 4Cii). The m^th^ columns of the firing-rate maps from different units were stacked to construct the population firing-rate matrix of the m^th^ parallel band (Figure 4Ciii). For the normalized correlograms, the firing-rate vectors of individual units were divided by the maximum firing rates before being stacked.

The correlograms were composed of the averaged correlation between pairs of population firing-rate matrices. The Pearson correlation was calculated between the same columns from the i^th^ and j^th^ population firing-rate matrices, and the correlations from each column were averaged and became the (i^th^, j^th^) element of the correlogram (Figure 4Ciii). Each bin of the correlograms represented the averaged correlation between two parallel band areas.

#### Place field detection

For each cell-session collected in the simple board or the plain board sessions that passed the spatial cell filter (see the *Spatial cell filtering* section), place field edges were detected for an area extending 30 cm beyond each side of the experiment board. These detection areas were 1.6 x 1.6 m for the simple boards and 1.7 x 1.7 m for the plain board. To construct the place fields, each area was partitioned into a matrix of 2 x 2 cm spatial bins, and the smoothed average firing rate for each bin was calculated as described in the *Firing rate map construction* section.

We binarized the firing-rate maps with floor-thresholds that were independently calculated for each cell-session. The spatial bins with firing rates larger than the thresholds were selected, and each connected group of bins was classified as a putative place field. The thresholds were set to maximize the differences between the mean firing rates within and outside of the place field(s), to minimize the variance of the firing rates outside of the place field(s), and to minimize the total size of the place field(s) (this last term was required to prevent all bins being included in the place field). In practice, we defined an error function of the threshold and optimized the error function with the *minimize_scalar* function under the *Scipy optimize* package [85] to find the best threshold θ that minimized the error function. The tolerance level was set at 10^-7^.

The error function was empirically defined as

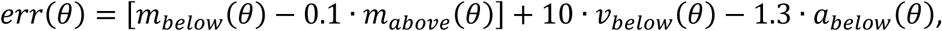

where *m*_*below*_(*θ*) was the mean firing rate of the bins with non-zero firing rates ≤ the current threshold *θ*, *m*_*above*_(*θ*) was the mean firing rate of the bins with firing rates > *θ*, *v*_*below*_(*θ*) was the firing rate variance of the bins with non-zero firing rates ≤ *θ*, and *a*_*below*_(*θ*) was the number of bins with non-zero firing rates ≤ *θ*, divided by the number of bins with non-zero firing rates. The search range for *θ* was limited to positive numbers ≤ the peak firing rate of the cell-session.

Since the same error function was used for different cell-sessions, some normalization for the means and variances of the firing rates across cell-sessions was necessary. For each cell-session, we first removed the firing rate outliers by calculating the quartiles and the interquartile range (*IQR*) of the non-zero firing rates, and truncated the firing-rate map at *Q*_1_ − *I*Q*R* and *Q*_3_ + *IQR*, where *Q*_1_ and *Q*_3_ were the first and third quartiles of the non-zero firing rates. For any element of the firing-rate map with non-zero value smaller than *Q*_1_ − *IQR* or larger than *Q*_3_ + *IQR*, the firing rate was re-assigned as *Q*_1_ − *IQR* or *Q*_3_ + *IQR*, respectively. We then normalized the truncated firing-rate map with the following rules: for the bins with non-zero firing rates, the truncated firing rates were transformed into standard scores, and the normalized firing rates were defined as the standard scores + 5; for the bins with zero firing rates, the normalized firing rates were still zero. The constant term (+5) was included to artificially differentiate the silent bins and the bins with non-zero firing rates. This preprocessing procedure was taken before optimizing the error function. Once the thresholds were determined based on the preprocessed firing-rate maps, the preprocessing procedure was reversed to recover the threshold, and the place fields were detected by binarizing the original firing-rate maps at the recovered threshold.

The connected bins with firing rates above threshold were grouped as putative place fields, and the contours of the putative place fields were then smoothed by opening and closing operations used in image processing [86]. An opening operator with a 3 x 3 bin square kernel (a 3 x 3 matrix with all 1s) was first applied to the putative fields to trowel small protrusions, followed by a closing operator with the same kernel to grout the small dents, and finished with a second opening operator with a cross-shaped kernel (a 4 x 4 matrix with 0s at the four corners and 1s at other locations) to eliminate any artificial protrusions created during the closing operation. After smoothing, the grouping of bins and putative field assignments were updated as above. Putative fields smaller than 35 bins, with peak firing rate < 0.1 Hz, or with no more than 30 in-field running spikes, were discarded, and the remaining qualified putative fields were labeled as place fields. For each spatial bin within a place field, we examined whether any of its adjacent bins (the bins above, below, to the left of, or to the right of the bin of interest) did not belong to the place field. If so the bin was labeled as belonging to the contour of the place field.

#### Boundary preference index (BPI) analyses

The spatial binning of the experiment boards and the construction of place field edges were described in the *Place field detection* section. For each simple board we examined whether spatial bins with high field edge occurrence were observed near the cue boundaries more often than expected by chance, by comparing the cumulative distribution functions (CDFs) of field edge occurrence. The differences between the area-under-curve (AUC) of the CDFs near and far away from the cue boundaries were computed and compared to the control data collected from the plain board.

To calculate the field edge occurrence, the simple boards were partitioned into the boundary zones and the non-boundary zones based on the distance to the boundary. The boundary zones were bands aligned with the cue boundaries, extended to the board edges and expanded 10 cm wide on each side of the cue boundaries (Figure 5A). For each spatial bin within the boundary or nonboundary zone and far from the board edges, the occurrence was defined as the number of fields with edges that overlapped with the bin. The zone boundaries were defined by linear equations, and a spatial bin could thus partially belong to the boundary and nonboundary zones simultaneously. For the spatial bin segregated by the zone boundaries, the bin would be assigned to the zone containing the larger proportion of the bin area. Since the place fields would be forced to end near the board edges, we excluded any spatial bin with center less than 10 cm from any of the board edges to avoid including place field edges that were not meaningful contributors to the analyses of surface cue boundary effects.

We calculated the boundary preference index (BPI) *a_b_* − *a_nb_* for each *simple board*, where *a_b_* was the AUC of the field edge occurrences within the boundary zone and *a_nb_* was the AUC within the nonboundary zone. The chance level of the AUC difference was determined by projecting the boundary and nonboundary zone demarcations from the simple boards onto the plain board and calculating the AUC difference of the data collected from the plain board accordingly. For all four simple boards, the same plain board and the same set of data recordings were used to calculate the control BPI. The plain board (1.1 x 1.1 m) was slightly larger than the simple boards (1 x 1 m), and therefore we rescaled the plain board data to fit the sizes of the simple boards.

To test whether the observed BPI was significantly higher than the control BPI, we separately and independently bootstrapped the place fields collected from the simple boards and the plain board 1,000 times. Each time N place fields were randomly resampled from the specified board with replacement, where N was the number of actual place fields collected from the board. The BPIs were calculated, and the difference between the observed and the control BPI (observed - control) was recorded in each trial. Based on the results obtained from the double rotation and the complex board data, we designed the simple board foraging task with the *a priori* prediction that field edges would concentrate near the cue boundaries, thus producing an observed BPI larger than the control BPI. The statistical significance was thus obtained by examining whether 95% of the bootstrapped BPIs was greater than 0 (i.e., a one-tailed test).

We also performed a permutation test to examine whether the BPI difference was significant. For each trial, the source labels of the place fields were shuffled and the fields were randomly reassigned to the simple board or the plain board. The BPI difference was calculated based on the shuffled field labels and the same process repeated 1,000 times. The observed BPI was considered significantly larger than the control BPI if the observed BPI difference was larger than or equal to the 1.25 percentile (significance level α = 0.05, one-tailed and Bonferroni corrected for 4 comparisons) of the shuffled distribution of the BPI difference.

#### Population vector direction of change analyses

For each location, we sought to determine which direction produced the maximum change in the population vector (PV) of firing rates between neighboring locations. The experiment boards were binned and the smoothed mean firing rate at each spatial bin was calculated as described in the *Firing-rate map construction* section. Since the binned rate maps allow calculation of movement angles in only 8 directions, none of which necessarily corresponded to the angle of the cue boundary, interpolation of the PV difference at arbitrary directions was necessary (Figure S6). The PV representing an arbitrary location (not restricted by the empirical binning) can be depicted by a continuous multivariate function *r*(*x*, *y*) which can be complex and implicit. Nevertheless, the tangent plane of *r*(*x*, *y*) can be estimated based on the binned rate maps even though *r*(*x*, *y*) itself is unknown. Taking each spatial bin as the reference point, we linearly approximated *r*(*x*, *y*) by its tangent plane *T*(*x*, *y*) and calculated the change of PV from the reference point to any neighboring locations. We quantified the difference between two PVs by the Euclidean distance between them and computed the direction *ω* with the largest PV difference (see Appendix for mathematical derivation of this procedure).

There are some noteworthy implicit rules applied to the searching of *ω* based on the linearity of the tangent plane. First, if the maximum PV difference was perceived at direction *ω*, the same amount of change would be observed at direction *ω*+180°. We therefore restricted *ω* to range from 0° to 180° while theoretically *ω* can range from 0° to 360° (0° is defined as parallel with the cue boundary for simplicity). Second, if the PV difference is only observed in the x (y) direction on the empirical binned rate map, *ω* would be the x (y) direction; if the PV difference is also observed in the y (x) direction, *ω* diverges from the x (y) direction. That is, the direction with the largest change would be perpendicular to the direction with the minimum change.

After the direction *ω* was calculated for each spatial bin, the direction vectors (unit vector with angle *ω*) within the boundary zone (or nonboundary zone) were concatenated and the length of the resulting vector was divided by the cell number to construct the mean direction vector of the boundary zone (or nonboundary zone). The mean vector length was then defined as the length of the mean direction vector.

## Acknowledgements

We thank Geeta Rao and Ravikrishnan Jayakumar for insightful comments on the manuscript. We thank Inah Lee, Yoganarasimha Doreswamy, Eric Hargreaves, Joshua Neunuebel, Sachin Deshmukh and Heekyung Lee for use of their double rotation data; and we thank Manu Madhav, Ravikrishnan Jayakumar, Vyash Puliyadi and Cheng Wang for assistance in experimental procedures and for technical support. This study is funded by U. S. Public Health Service grants R01 NS09456 and R01 MH094146.

## Author Contributions

Conceptualization, C.W. and J.J.K.; Methodology, C.W. and J.D.M.; Formal analysis, C.W.; Investigation, C.W.; Writing – Original Draft, C.W. and J.J.K.; Writing – Review & Editing, C.W., J.D.M., and J.J.K.; Supervision, J.J.K.; Funding Acquisition, J.J.K.

## Declaration of Interests

The authors declare no competing interests.

## Appendix

The firing rates of different place cells formed the population vector of firing rate *r⃑*, defined as a function of the physical location of the animal,

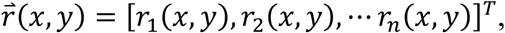

where *r*_*i*_(*x*, *y*) is the smoothed mean firing rate of the *i*^th^ place cell. When the animal moves from (*x*_0_, *y*_0_) to (*x*_1_, *y*_1_) on the experiment board, there is a corresponding change in the population vector,

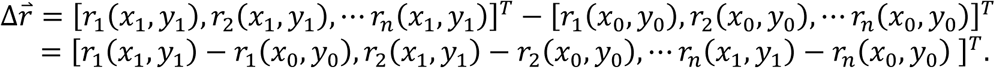

For any point (*x*, *y*) close to (*x*_0_, *y*_0_), *r⃑*(*x*, *y*) can be linearly approximated by its tangent plane *T*(*x*, *y*) at (*x*_0_, *y*_0_). Thus, Δ*r⃑* can be approximated by the corresponding change on the tangent plane:

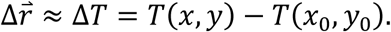

In order to compute *T*(*x*, *y*), for each spatial bin we calculated the Jacobian matrix,

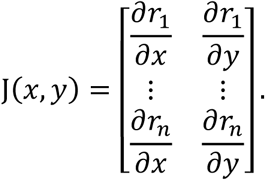

The Jacobian matrix at (*x*_0_, *y*_0_) transforms a point close to (*x*_0_, *y*_0_) to its corresponding location on the tangent plane *T*(*x*, *y*). In other words, we can linearly approximate *r⃑*(*x*, *y*) near (*x*_0_, *y*_0_) by using the Jacobian matrix *J*(*x*_0_, *y*0) to calculate the tangent plane,

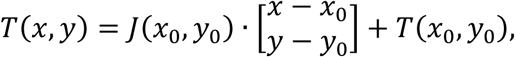

and, therefore, the change in population firing rate vectors,

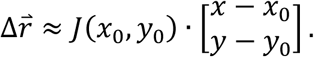

We quantified the magnitude of Δ*r⃑* as its Euclidean vector norm, ‖Δ*r⃑*‖. The heading direction on the x-y plane that produces the largest change in the population vector of firing rates would thus also maximize displacement along the tangent plane. In other words, we searched for the heading direction [*x* − *x*_0_, *y* − *y*_0_]*T* = [cos *ω*, sin *ω*]*T* that maximized ‖Δ*r⃑*‖, i.e., 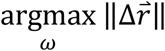, by equivalently solving

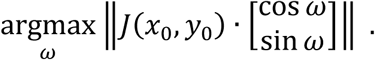

The maximal heading *ω* may theoretically range from 0° to 360°. However, the linearity of function Δ*T* and the symmetry of the vector norm combine to produce equivalent rate changes for shifts of 180°,

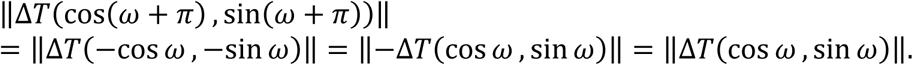

Thus, we restricted solutions for the maximal heading *ω* to the range [0°, 180°].

